# Disruption of R2TP-mediated PIKK maturation with *de novo* designed binders

**DOI:** 10.64898/2026.10.26.761954

**Authors:** Nayim González-Rodríguez, Andrés López-Perrote, Clara Reglero, Jaime Alegrio-Louro, Marta Failde-Soler, Carlos Chacón-Sánchez, Elena Blanco-Romero, Ester Casajús-Pelegay, Gema López-Pernas, Ivó H. Hernández, Matilde Murga, Oscar Fernandez-Capetillo, Oscar Llorca, Rafael Fernandez-Leiro

**Author notes:** Co-corresponding authors: Oscar Llorca and Rafael Fernandez-Leiro. These authors contributed equally (Clara Reglero, Jaime Alegrio-Louro, Marta Failde-Soler). Hull York Medical School, Department of Biology, York Biomedical Research Institute, University of York, York, UK.

## Abstract

Phosphatidylinositol 3-kinase–related kinases (PIKKs), including ATM, ATR, DNA-PKcs, SMG1, TRRAP and mTOR, are regulators of gene expression, DNA damage response and metabolism. Their maturation depends on R2TP, a co-chaperone complex comprising the RUVBL1/2 AAA+ ATPase, RPAP3 and PIH1D1. While small-molecule RUVBL1/2 inhibitors have shown promise in cancer models, the presence of RUVBL1/2 in other complexes raises concerns about specificity. Here, we identified the RUVBL1/2-RPAP3 interface as a vulnerability, exploiting it using *de novo* designed protein binders that occupy the RPAP3-binding site. These subnanomolar-affinity binders dismantle R2TP, depleting levels of all PIKKs while sparing other RUVBL1/2-containing complexes, including chromatin remodelers INO80, SRCAP and TIP60. CryoEM, proteomic, biochemical and cellular analyses revealed that these binders outcompete RPAP3, trapping RUVBL1/2 in a catalytically inactive conformation. Our findings establish selective disruption of R2TP by artificial proteins as a strategy to control PIKK abundance.

## 1. INTRODUCTION

**P**hosphatidylinositol 3-kinase–related kinases (PIKKs) constitute a family of large proteins that regulate diverse cellular pathways and have emerged as interesting therapeutic targets, particularly in cancer. The PIKK family comprises kinases like mTOR, a central regulator of cell growth, metabolism, and nutrient sensing; ATM, ATR and DNA-PKcs, which participate in DNA damage signaling and repair; SMG1, which regulates nonsense-mediated mRNA decay (NMD); and the pseudokinase TRRAP, which functions in transcriptional regulation^1^. PIKKs share conserved structural features, and they function within large multiprotein assemblies. mTOR, for example, is only active as part of the larger multi-subunit complexes mTOR complex 1 (mTORC1) and 2 (mTORC2)^2^.

The large size and modular architecture of PIKKs pose significant challenges for their folding, stability and assembly into functional multiprotein complexes. Their maturation critically depends on specialized chaperone systems including the RUVBL1/2-RPAP3-PIH1D1 (R2TP) and TELO2-TTI1-TTI2 (TTT) complexes^3–5^ acting in concert with the HSP90 chaperone^6–8^. Although the molecular mechanisms underlying this pathway remain incompletely understood, current evidence suggests that TTT associates co-translationally with nascent PIKKs to promote their folding^9,10^. TTT transfers partially folded clients to the R2TP-HSP90 machinery for further maturation and assembly into functional higher-order complexes through interactions with R2TP^6–9^ and, possibly, additional factors such as GNB1L^11^. Accordingly, depletion or mutation of TTT or R2TP components, as well as pharmacological inhibition of HSP90, results in reduced levels of all PIKKs^3–5^. RUVBL1/2 is an AAA+-ATPase complex, and mutants lacking ATPase activity also result in reduced levels of PIKKs, indicating that its ATPase activity is also required for R2TP-dependent PIKK maturation^4^. Likewise, the assembly of at least the ATR-ATRIP and the mTORC1 and mTORC2 complexes depends on R2TP^3,5^. Therefore, the R2TP complex acts as a central regulator of PIKK maturation and several lines of evidence support the therapeutic potential of targeting R2TP and RUVBL1/2. Indeed, CB-6644, a selective allosteric RUVBL1/2 ATPase inhibitor, has shown activity in multiple cancer models^12^, and related compounds showed efficacy in non-small cell lung cancer and enhanced radiosensitivity in prostate cancer^13–16^. Interestingly, a phenotypic screen searching for compounds that induce cell death selectively in cancer cells with high dependence on mTORC1 identified a compound that disrupts the RUVBL1/2-TTT interaction^17^.

RUVBL1/2 is arranged as a heterohexameric ring that presents two distinct faces: the AAA+-face, containing its catalytic sites, and opposed to it, the DII-domain face, involved in interactions with diverse partners. RUVBL1/2 participates in multiple cellular processes beyond R2TP^6,18^, such as the Fanconi anemia pathway^19^, microtubule nucleation^20^, or MYC-dependent pathways in neuroblastoma cells^13^. Notably, RUVBL1/2 is a core component of several chromatin remodeling complexes, including INO80, SRCAP, and TIP60, where it engages additional subunits through its DII-domain face^21–23^. Consequently, ATPase inhibition may broadly perturb other cellular processes. This problem has already been recognized for other AAA+-ATPases, where it has been proposed that disrupting specific protein-protein interactions, rather than their enzymatic activity, could achieve greater selectivity^24^.

We therefore decided to selectively disrupt R2TP by targeting the interaction between RUVBL1/2 and RPAP3. This strategy exploits three structural features of the complex. First, R2TP integrity relies exclusively on the interaction between the RPAP3 RUVBL-binding domain (RBD) and RUVBL2, essential to recruit all other components of R2TP and HSP90 to RUVBL1/2^25,26^. Second, the RBD binds to the AAA+-face of RUVBL1/2, spatially separated from the DII-face that mediates protein interactions within most of the other RUVBL1/2-containing complexes^25^. Third, structural studies revealed that RU-VBL1/2 adopts an ADP-bound conformation in the R2TP complex that differs from those observed in other RUVBL1/2 assemblies^27^, enabling conformation-selective inhibition. Therefore, we designed *de novo* high-affinity, RUVBL1/2-specific binders (RBs) against this interface. Using cryo-electron microscopy (cryoEM), proteomics, biochemical, and cellular analyses, we show that these binders selectively disrupt R2TP and reduce cellular PIKK abundance as a result. Thus, in this work, we establish a strategy to selectively perturb the R2TP-dependent PIKK maturation pathway while sparing other cellular RUVBL1/2 functions.

## 2. RESULTS

### 2.1. Iterative design and refinement yield subnanomolar RU-VBL1/2 binders

R2TP formation depends on the interaction between RPAP3 and the hexameric RUVBL1/2 ring. The C-terminal RBD in RPAP3 recognizes RUVBL2^25,26^ and disrupting this interaction is sufficient to disengage RPAP3-PIH1D1 from RUVBL1/2^26,28^. We aimed to prevent PIKK maturation by affecting R2TP assembly through the design of binders that occupy the RBD binding site on RUVBL1/2, preventing RPAP3 binding through direct competition. We hypothesized that high-affinity RBs could not only prevent RPAP3 binding but also disrupt existing R2TP complexes (**Fig. 1a**). We targeted residues on RUVBL2 essential for RBD binding, V423, I426 and Y430, that interact with RPAP3 residues M626 and F630^26,28^ (**Fig. 1b**). Given that the RBD interacts with RUVBL1/2 with high affinity (*K*_*D*_ ≈ 4 nM)^26^, we reasoned that effective competition would require even higher-affinity RBs.

**Figure 1.**
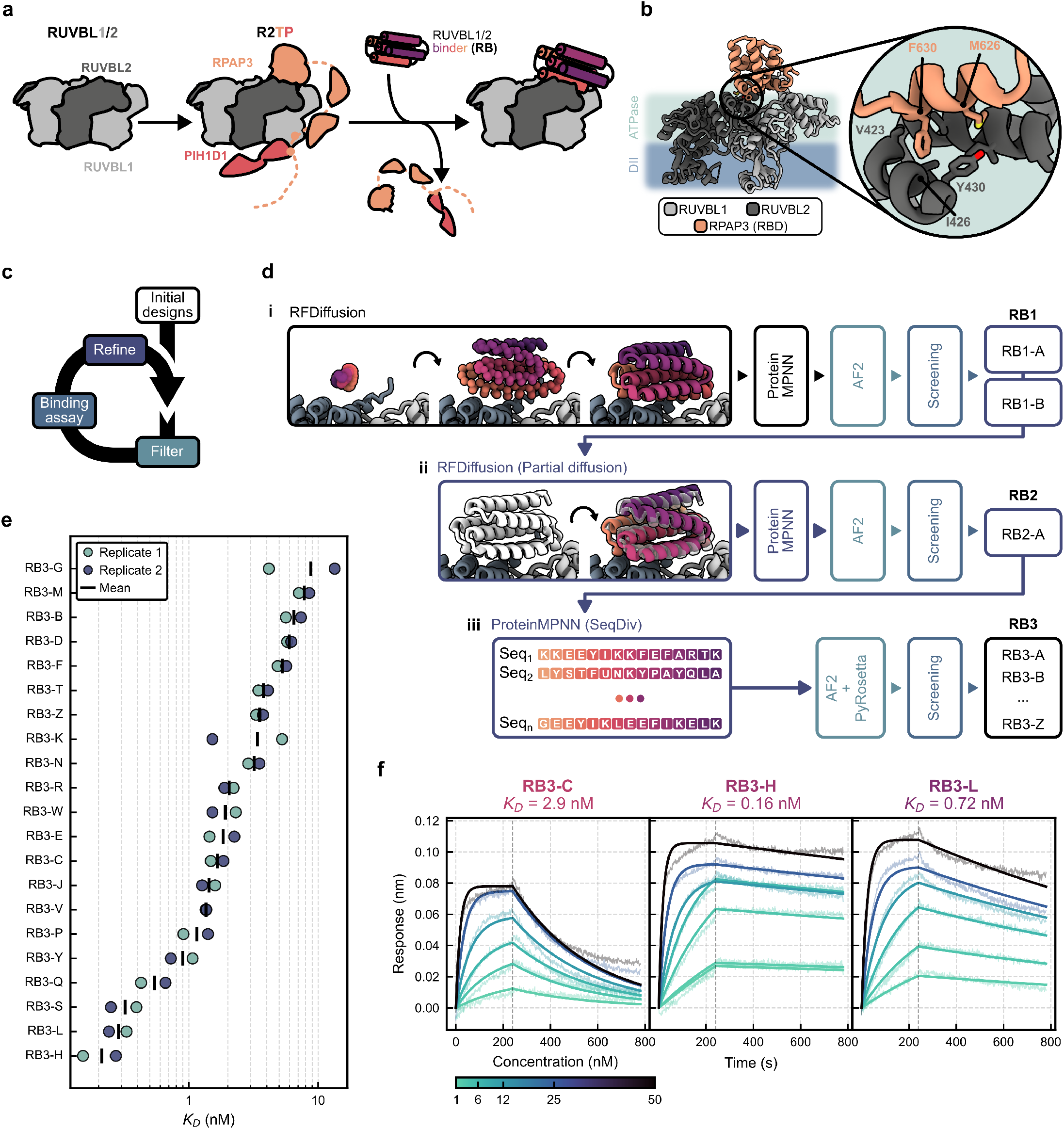
*De novo* design and affinity characterization of RBs. **a**, RUVBL1/2 recruits RPAP3 and PIH1D1 to assemble R2TP. By designing binders *de novo*, we aim to prevent the binding of and/or outcompete RPAP3 from its RUVBL1/2 binding site, thus preventing R2TP assembly. **b**, Cartoon representation of the RPAP3 binding domain in RUVBL2, highlighting the hydrophobic pocket buried on its ATPase face where RPAP3 inserts residues F630 and M626. **c**, Iterative binder design strategy. To avoid costly large-scale experimental screening, we aimed to perform *de novo* design iteratively. Initial designs were strictly filtered *in silico*, and a limited number of candidates was tested experimentally. Initial binders underwent successive rounds of *in silico* optimization to improve their affinities. **d**, Schematic representations of the strategy followed during (i) the first design iteration, RFDiffusion (RFD) was used to generate backbones *de novo* and ProteinMPNN for inverse folding; (ii) Partial diffusion refinement of binders. RFD was used to add noise to the positions of RB1-A and -B backbones to be denoised into refined backbones, and ProteinMPNN again used for inverse folding; (iii) Sequence Diversity refinement of RB2s. AF2 was used for scoring all designs across all design stages. **e**, Sorted plot of all *K*_*D*_ values from BLI binding characterization of RB3s. Vertical black lines represent average *K*_*D*_ values (n=2). **f**, BLI characterization of the RB3s selected for further characterization, RB3-C, -H and -L.

The lack of *bona fide* metrics that predict binding of *de novo* designed binders implies the need for large-scale computational design campaigns followed by costly experimental validation^29^. We reduced the computational design burden by using BinderFlow, a computational pipeline we developed to run long unsupervised design campaigns^30^. To minimize the number of DNA fragments synthesized and experimental screening, we adopted an iterative design strategy. It consisted of (1) moderately sized design campaigns, (2) stringent *in silico* filtering, (3) validation of a limited set of candidates, and (4) feeding validated RBs back into rounds of computational refinement, iteratively improving affinity (**Fig. 1c**).

We performed an initial round of design using the standard BinderFlow pipeline. We used RoseTTAFold Diffusion (RFDiffusion) to compute ∼12,000 backbones *de novo*, ranging from 60 to 220 residues in length, and inversely folded each of them with ProteinMPNN^31^. AlphaFold2 (AF2)^32,33^ was used to predict the structure of each binder in complex with RUVBL1/2 to filter candidates *in silico* as previously described^34^ (**Fig. 1d, i**). After filtering by pLDDT, iPAE and visual inspection to ensure a sufficiently large hydrophobic core, five designs, RUVBL1/2 binders 1 (RB1), were selected as candidates (**Supplementary Fig. S1a, b**). Their coding sequences were synthesized as DNA fragments and cloned individually in pRSF-Duet vectors, overexpressed in *Escherichia coli* and purified. Binding to RUVBL1/2 was initially assessed by pull-down and immunoblotting. RB1-A and RB1-B showed detectable binding to purified RUVBL1/2 (**Supplementary Fig. S1c**), corroborated by their coelution upon size exclusion chromatography (**Supplementary Fig. S1d**). We used isothermal titration calorimetry (ITC) to determine the affinity of their interaction with RUVBL1/2, obtaining *K*_*D*_ = 1.4 *μ*M and *K*_*D*_ = 6.3 *μ*M for RB1-A and RB1-B, respectively (**Supplementary Fig. S1e**).

Both RBs were subjected to a second round of design to optimize backbone geometry, aiming to improve their affinity. Using Partial Diffusion^35^, the C_*α*_ coordinates of RB1 backbones were perturbed by small amounts of gaussian noise and re-diffused to refine shape complementarity with targeted RUVBL1/2 (**Fig. 1d, ii**). Top-ranked designs from each campaign, assessed by iPAE, were selected (**Supplementary Fig. S2a**), purified and characterized by ITC. RB2-A bound RUVBL1/2 with *K*_*D*_ = 0.19 *μ*M, a ∼10× increase in binding affinity relative to its parental design RB1-A (**Supplementary Fig. S2b**). The affinity increase is entropy-driven, likely due to the improved shape complementarity between the redesigned backbone in RB2-A and RUVBL1/2 (**Supplementary Fig. S2c**).

Given that RB2-A still presented lower affinity for RUVBL1/2 than RPAP3^26^, we subjected this design to a final round of computational refinement. We used BinderFlow’s implementation of the Sequence Diversity routine, in which thousands of ProteinMPNN sequences per backbone are sampled and predicted^30^ (**Fig. 1d, iii**). The high *in silico* success rate of this campaign allowed us to enforce stringent AF2 and biophysical filters to select candidate designs (**Supplementary Fig. S2d**). This time, we expanded the scale of the experimental validation campaign to maximize the likelihood of finding high-affinity interactors, and 26 candidates were successfully overexpressed and purified (RB3-A through RB3-Z). We assessed their binding to RUVBL1/2 by pull-down, in which untagged RUVBL1/2 complexes were flowed through Ni^2+^-NTA beads pre-incubated with His-tagged RB3s. All overexpressed RB3s coprecipitated RUVBL1/2 (**Supplementary Fig. S2e**). We characterized their binding affinities by biolayer interferometry (BLI), measuring their interaction to immobilized, biotinylated RUVBL1/2. The resulting affinities spanned over three orders of magnitude, ranging from *K*_*D*_ ∼ 10 nM (RB3-G) to *K*_*D*_ ∼ 0.2 nM (RB3-H), despite relatively similar sequences (**Fig. 1e, Supplementary Fig. S3**). We chose three of the highest-affinity interactors, RB3-C, -H and -L, for further characterization and determined their affinities with higher precision by BLI, resulting in *K*_*D*_ = 2.9, 0.16 and 0.72 nM, respectively (**Fig. 1f, Supplementary Fig. S3**).

In summary, we designed *de novo* RUVBL1/2 binders with sub-nanomolar affinity by combining small-scale experimental screening with iterative rounds of computational refinement. We achieved a ∼10000× improvement in binding affinity from the initial design RB1-A (*K*_*D*_ = 1.4 *μ*M) to the optimized binder RB3-H (*K*_*D*_ = 0.16 nM), exceeding that of RPAP3 by ∼25-fold (*K*_*D*_ ∼ 4 nM)^26^.

### 2.2. Structural characterization of RUVBL1/2-RB complexes

AF2 predictions of all RB3s share a 6-*α*-helix bundle topology inherited from the parental design RB1-A (**Fig. 2a**), despite showing drastically higher affinity. In all cases, helices *α*1, *α*2 and *α*4 are predicted as oriented towards RUVBL1/2, with *α*2 buried across the RBD-binding surface. To confirm whether RBs bound specifically to the RBD-interaction surface, we determined experimental structures of some of the highest-affinity RBs: RB3-C, -H and -L. In addition, we also determined the structures of their parental designs, RB1-A and RB2-A, to understand the structural determinants of the increase in affinity across design generations.

**Figure 2.**
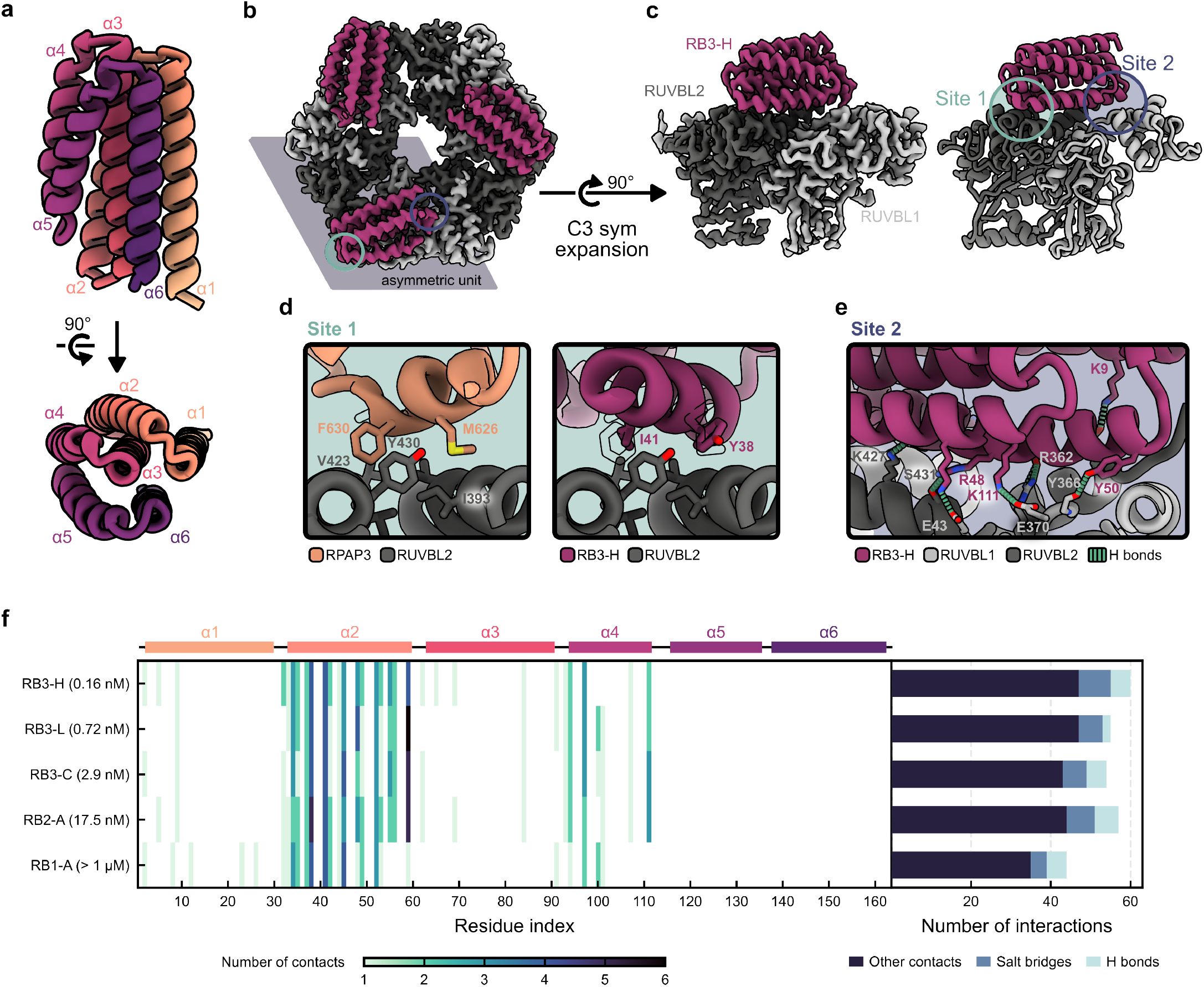
CryoEM and structural characterization of RUVBL1/2-RB complexes. **a**, Cartoon representation of RB3-H as a representative example of the binders *α*-helical bundle fold, showing two orthogonal views. **b**, Surface representation of the raw, refined map of the RUVBL1/2-RB3-H C3-symmetric complex. RB3-H colored in purple, RUVBL1/2 in shades of gray. **c**, Surface representation of the raw, refined map of the RUVBL1/2-RB3-H complex asymmetric unit after symmetry expansion (left), and cartoon representation of the corresponding atomic model (right). RB3s recognize RUVBL1/2 through two sites. **d**, Structural details of the RPAP3-RUVBL1/2 interaction in Site 1 (left) and equivalent view of the RUVBL1/2-RB3-H interaction, with RPAP3 M626 and F630 represented as outlines (right). **e**, Structural details of the RUVBL1/2-RB3-H interaction in Site 2. **f**, Heatmap of per-residue contacts between binders and RUVBL1/2 with the RB secondary structure indicated schematically on top of the heatmap (left) and bar plot of the number of total interactions (right).

Each binder was incubated with RUVBL1/2 and vitrified for cryoEM data collection (**Supplementary Table S1**). Upon 2D classification of the extracted particles, most class averages corresponded to side projections of dodecameric RUVBL1/2 complexes, consistent with previous observations^25^ (**Supplementary Fig. S4a**). Our goal was to characterize the RUVBL1/2-RB interaction surface. Therefore, we reconstructed only one hexameric ring, avoiding the flexibility of the dodecamer interface^36–38^, and obtained high-resolution reconstructions for all five complexes. We observed helical densities consistent with the RB 6-*α*-helix fold bound to each RUVBL2 subunit at the AAA+-face of the RUVBL1/2 ring. In R2TP, each of the three RUVBL2 subunits can be recognized by one RBD^25^, and in our RUVBL1/2-RB complexes each of these sites was also occupied by one RB (**Fig. 2b, Supplementary Fig. S4b-d**). Uneven RB density quality across binding sites, particularly obvious in RB1-A and RB2-A, suggested partial occupancy of RBD binding sites, as shown by local scale analyses^39^ (**Supplementary Fig. S4e**). We exploited the C3 symmetry of the complex by applying symmetry expansion and focused 3D classification without alignment to remove RUVBL1/2 dimers lacking bound RBs (**Fig. 2c, Supplementary Fig. S5a**,**b**). The improved quality of the RB densities allowed confident model building at the RUVBL1/2-RB interface (**Supplementary Fig. S5c-f**).

All RBs featured the 6-*α*-helix topology predicted by AF2 with remarkable accuracy, with all-atom RMSD values below 0.7 Å (**Supplementary Fig. S6a**,**b)**. All RBs occupied the intended RBD-binding site on RUVBL1/2, and interacted mainly through two distinct sites, one distal to the hexamer center, Site 1, and another proximal one, Site 2 (**Fig. 2b,c**). The interactions at Site 1 occlude the hydrophobic pocket where the F630 and M626 residues from RPAP3 are inserted, key for the RBD-RUVBL1/2 interaction^26,28^. These are replaced by Y38 and I41, conserved in all characterized RBs across design generations, indicating that these positions are also particularly relevant for RUVBL1/2-RB high-affinity interactions (**Fig. 2d, Supplementary Fig. S6c**,**d**). In contrast to RBD, which contacts only at Site 1, RBs establish an extensive network of additional interactions at Site 2, contacting both RUVBL1 and RUVBL2 (**Fig. 2e**), which explains the potential of RBs to achieve higher affinity than RPAP3. The optimized geometry of RB2 and RB3 backbones allows for positioning *α*4 closer to RUVBL1/2, enabling additional polar contacts (**Supplementary Fig. S6d**). The increased affinity of refined binders in comparison with RB1-A can thus be explained by the optimized backbone and sequences, which better complement the targeted surface shape and allow establishing a new set of interactions involving the C-ter end of

In summary, our cryoEM analyses demonstrate that RBs successfully target the intended RBD-binding surface of RUVBL1/2, occupying the main RPAP3-binding pocket. By mimicking the hydrophobic interactions required for RPAP3 binding and extending its interface with a new set of contacts, they achieve affinities exceeding those of the natural ligand by over an order of magnitude.

### 2.3. RBs specifically outcompete RPAP3 from binding RUVBL1/2

RBs were designed to occupy the RPAP3 RBD-binding site and bind RUVBL1/2 with higher affinity than RPAP3, thus we predicted that they should efficiently outcompete the RUVBL1/2-RPAP3 interaction. To test this, we reconstituted a minimal R2TP complex consisting of RUVBL1/2 bound to a Strep-tagged RBD from RPAP3. The purified complex was immobilized on StrepTactin resin and incubated with increasing concentrations of RB1-A, RB2-A and RB3-H (**Fig. 3a**). Displacement of RUVBL1/2 from immobilized RBD was monitored by SDS-PAGE. All three binders displaced RUVBL1/2 in a concentration-dependent fashion, even at substoichiometric ratios. Competition efficiency increased across design generations, with higher-affinity RB requiring progressively lower concentrations to disrupt preformed RUVBL1/2-RBD complexes (**Fig. 3b**). These results demonstrate that the RUVBL1/2 binders can effectively compete with RPAP3 for the RBD-binding site *in vitro* disrupting pre-assembled R2TP, and confirm the prediction that effective competition benefits from high-affinity binding.

**Figure 3.**
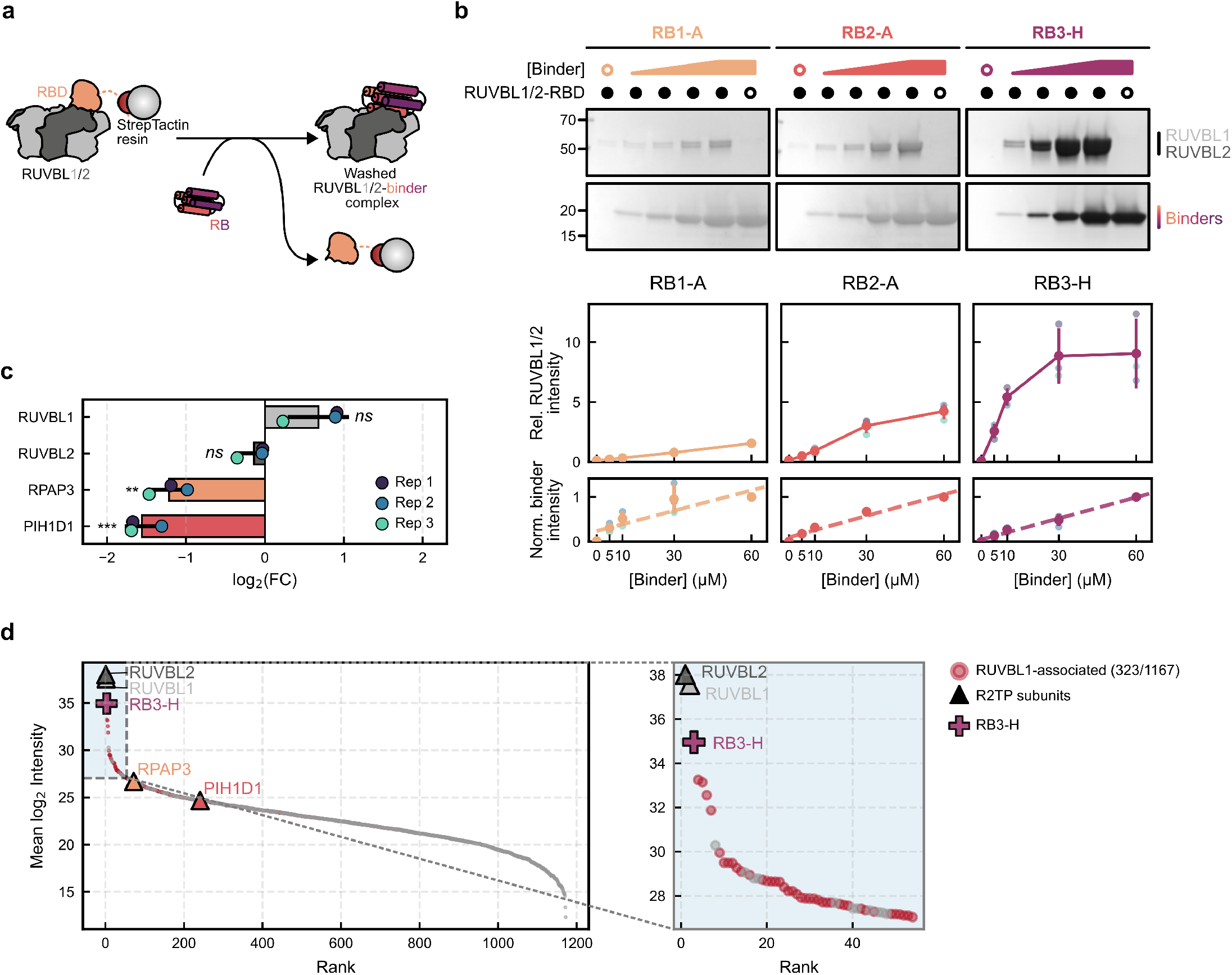
RBs disrupt R2TP assembly *in vitro* and in cells. **a**, Schematic representation of the competition assay between purified RUVBL1/2-RBD complexes and RBs. Pre-assembled RUVBL1/2-RBD complexes bound to StrepTactin resin are washed with RBs in increasing concentrations and disruption is assessed by the amount of RUVBL1/2 displaced. **b**, SDS-PAGE analyses of wash fractions from the competition assay depicted in panel a (top) and band intensity quantification (n = 3, each batch of gels processed in parallel) (bottom). **c**, Effect of RB3-H overexpression on R2TP assembly. Endogenous RUVBL1 was pulled down from HEK293T whole cell lysates transfected with RB3-H-containing or mock plasmids, and the eluates were analyzed by mass spectrometry to quantify the abundance of its partners. The bar plot represents the average and SD of differential abundance of R2TP subunits (n=3). Asterisks indicate p-values from moderated t-tests (limma) adjusted by Benjamini–Hochberg (FDR < 0.05). ns: p > 0.1, **: p < 0.01, ***: p < 0.001. **d**, Sorted intensity plot of RB3-H interactors (left), with the shaded light-blue region shown magnified (right). RB3-H was overexpressed and pulled down from HEK293T whole cell lysates, and the eluates were analyzed by mass spectrometry. Dots in red indicate proteins significantly detected in endogenous RUVBL1 pull-downs, indicating indirect interaction through RUVBL1, rather than off-target binding of RB3-H.

To test whether RBs can also outcompete RPAP3 in a cellular context, we immunoprecipitated endogenous RUVBL1 from HEK293T cells overexpressing RB3-H and compared the abundance of co-immunoprecipitated R2TP components to that of transfection controls by mass spectrometry. Upon RB3-H overexpression, co-precipitated RPAP3 and PIH1D1 levels significantly decreased by ∼60% and ∼70%, respectively. In contrast, the abundance of RUVBL2 and other RUVBL1/2 associated factors, such as the TTT complex or HSP90, did not change significantly (**Fig. 3c, Supplementary Fig. S7a**). These results indicate that RB3-H efficiently outcompetes RPAP3 from its binding to RUVBL1/2 in a cellular context, preventing RPAP3-PIH1D1 recruitment and disrupting R2TP assembly.

RFD-designed binders frequently target hydrophobic surfaces, with binding driven primarily by shape complementarity^40^. Because hydrophobic interfaces are ubiquitous in proteins, we next assessed the specificity of the highest-affinity binder, RB3-H. RB3-H was overexpressed in HEK293T cells, used as bait for pull-down and eluates were analyzed by mass spectrometry. As expected, RUVBL1 and RUVBL2 were the most abundant interactors. Notably, most of the remaining high-confidence interactors had also been detected in the endogenous RUVBL1 immunoprecipitation, strongly suggesting that they are RUVBL1/2 partners, co-eluting with the RUVBL1/2-RB complex, rather than non-specific RB3-H interactors (**Fig. 3d, Supplementary Fig. S7b**).

Together, these results show that high-affinity RBs specifically target RUVBL1/2 in cells, effectively outcompete RPAP3, and dismantle and/or prevent the assembly of R2TP.

### 2.4. RBs are allosteric inhibitors of RUVBL1/2

ATP binding and hydrolysis are coupled to conformational changes in the RUVBL1/2 ATPase ring that modulate its interaction with other proteins^41^. Conversely, interaction with certain clients can also modulate ATP hydrolysis^27,28,41^. We therefore asked whether RB binding affected RUVBL1/2 ATPase activity. Addition of either full-length RPAP3 or isolated RBD reduced ATP hydrolysis by ∼30%, and the low-affinity binder RB1-A had a similar effect. In contrast, RB2-A, RB3-C, -H and -L inhibited ATPase activity by ∼70-80% (**Fig. 4a**). Thus, RBs are allosteric RUVBL1/2 inhibitors, and while affinity alone does not fully explain ATPase inhibition (compare RB1-A and RBD), higher-affinity RBs seem to induce stronger ATPase inhibition.

**Figure 4.**
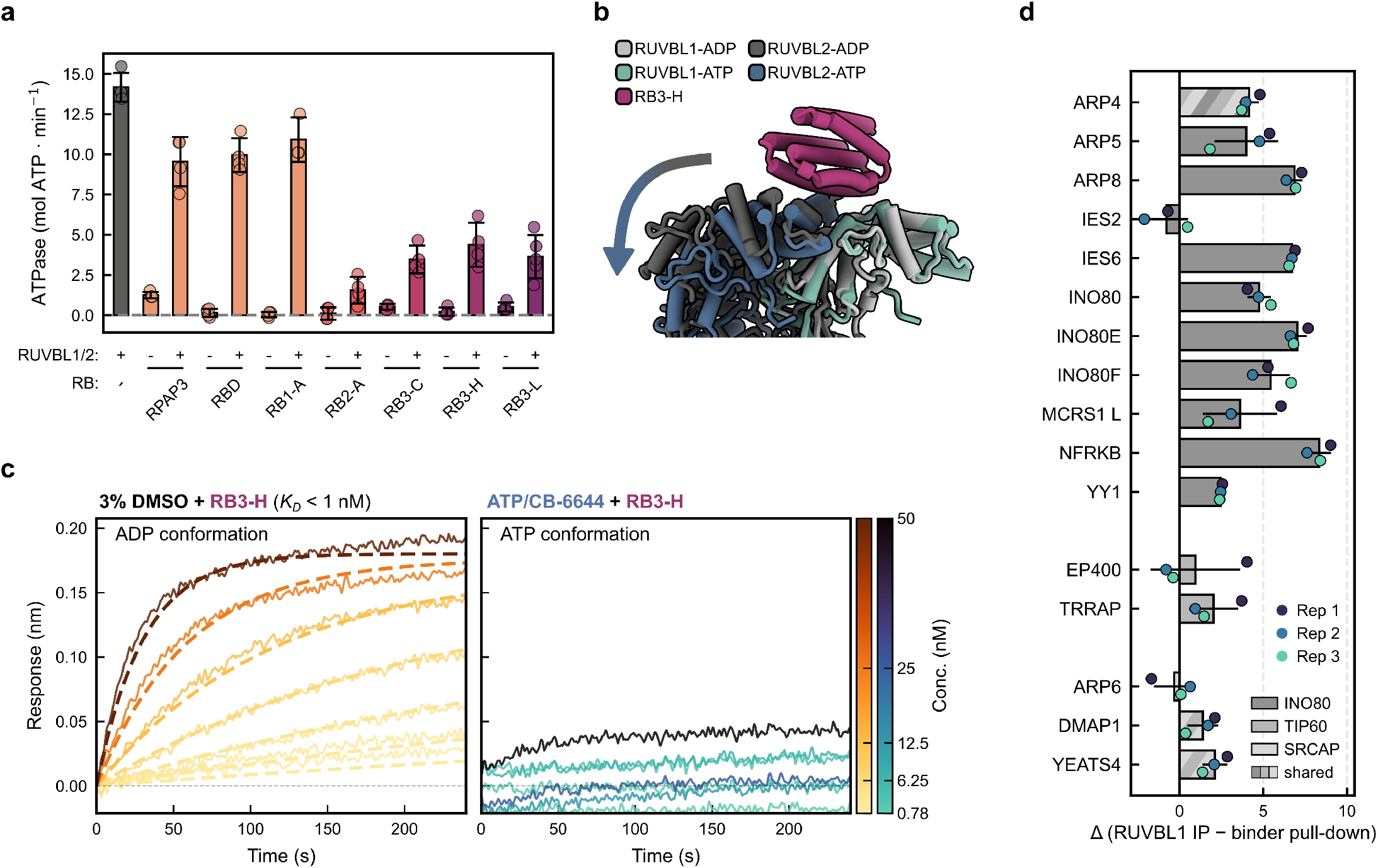
RBs allosterically inhibit RUVBL1/2 ATPase activity. **a**, ATP catalytic rates of purified RUVBL1/2 in presence of RPAP3, RBD and RBs. Bars represent average and SD; dots represent individual measurements (n=3-4). **b**, Comparison of the conformations of RUVBL1/2 ATPase face in ATP-bound (blue shades) and ADP-bound conformations (grey shades). **c**, BLI association traces for RB3-H binding to immobilized, biotinylated RUVBL1/2 complexes, in buffer supplemented with DMSO, rendering a RUVBL1/2 ADP-bound conformation (left), or ATP and CB-6644, rendering a trapped ATP-bound conformation (right). Dashed lines indicate globally fitted association kinetics. **d**, Stoichiometry of INO80, TIP60 and SRCAP subunits relative to RUVBL1, comparing an endogenous RUVBL1 immunoprecipitation with RB3-H pull-down. For each protein, log_2_(protein/RUVBL1) was computed per replicate. Δ = mean(RUVBL1 IP) - mean(binder pull-down). Δ > 0 means that, per unit RUVBL1, the protein co-purifies more with endogenous RUVBL1 than with the binder. Bars represent average Δ across three replicates, error bars, SD. Proteins absent from the dataset or detected in < 2 replicates of the RUVBL1 IP were excluded.

Notably, the ADP- and ATP-bound conformations of RUVBL1/2 differ substantially^27^ (**Supplementary Fig. S8a**). In the ADP-bound conformation, the ATPase face of the ring adopts a planar conformation that is competent for RPAP3 binding, whereas when bound to ATP, the ATPase face of RUVBL1/2 bends, remodeling the RUVBL2 RBD-binding surface and repositioning most RB-interacting residues (**Fig. 4b**). We designed RBs to target the ADP-bound RUVBL1/2 conformation, and this was confirmed by the cryoEM analyses (**Supplementary Fig. S5**). Together with the slow dissociation rates observed for the highest-affinity RBs (**Supplementary Fig. S3a**), these structures suggest that RB binding stabilizes the ADP-bound state, disfavoring the conformational changes required for ATP binding and hydrolysis, consistent with the inhibition observed.

To directly test this model, we asked whether RB3-H preferentially binds the ADP-bound conformation of RUVBL1/2. We measured RB3-H binding to immobilized RUVBL1/2 by BLI supplemented with ATP and CB-6644, an allosteric inhibitor shown to trap the ATP-bound conformation^27^. Under these conditions, RB3-H binding was dramatically reduced when compared to unsupplemented conditions (**Fig. 4c**), in which RUVBL1/2 is bound to ADP, as shown by our cryoEM analysis (**Supplementary Fig. S5f**). These data indicate that RB3-H displays a strong conformational preference for the ADP-bound state.

Such conformational bias should reduce RB binding to RUVBL1/2 assemblies that adopt the ATP-bound conformation. Consistent with our previous observations for the INO80 complex^27^, comparison of all available structures revealed that RUVBL1/2 adopts a similar ATP-bound-like conformation in the chromatin-remodeling complexes INO80, TIP60 and SRCAP^21,22,42^ (**Supplementary Fig. S8b**). Supporting this model, immunoprecipitation of RB3-H followed by mass spectrometry showed much lower amounts of chromatin remodeling complex subunits than immunoprecipitation of endogenous RUVBL1, normalized by the amount of precipitated RUVBL1 (**Fig. 4d**). Together, these results show that RBs allosterically inhibit RUVBL1/2 catalytic activity, likely stabilizing the ADP-bound conformation. This conformational selectivity implies that RBs are significantly less efficient in recognizing RUVBL1/2 in the ATP-bound conformation, providing a mechanistic explanation for the selective disruption of R2TP while sparing chromatin-remodeling complexes.

### 2.5. RBs reduce the abundance of all PIKKs

R2TP activity is necessary for the stability of all PIKKs^3–5^. Preventing the RUVBL1/2-RPAP3 interaction is therefore expected to reduce their cellular abundance. Indeed, transient overexpression of RB3-C, -H or -L significantly reduced the levels of ATR, mTOR and TRRAP in whole-cell lysates 72 h post-transfection in comparison with mock transfections (**Fig. 5a**,**b**).

**Figure 5.**
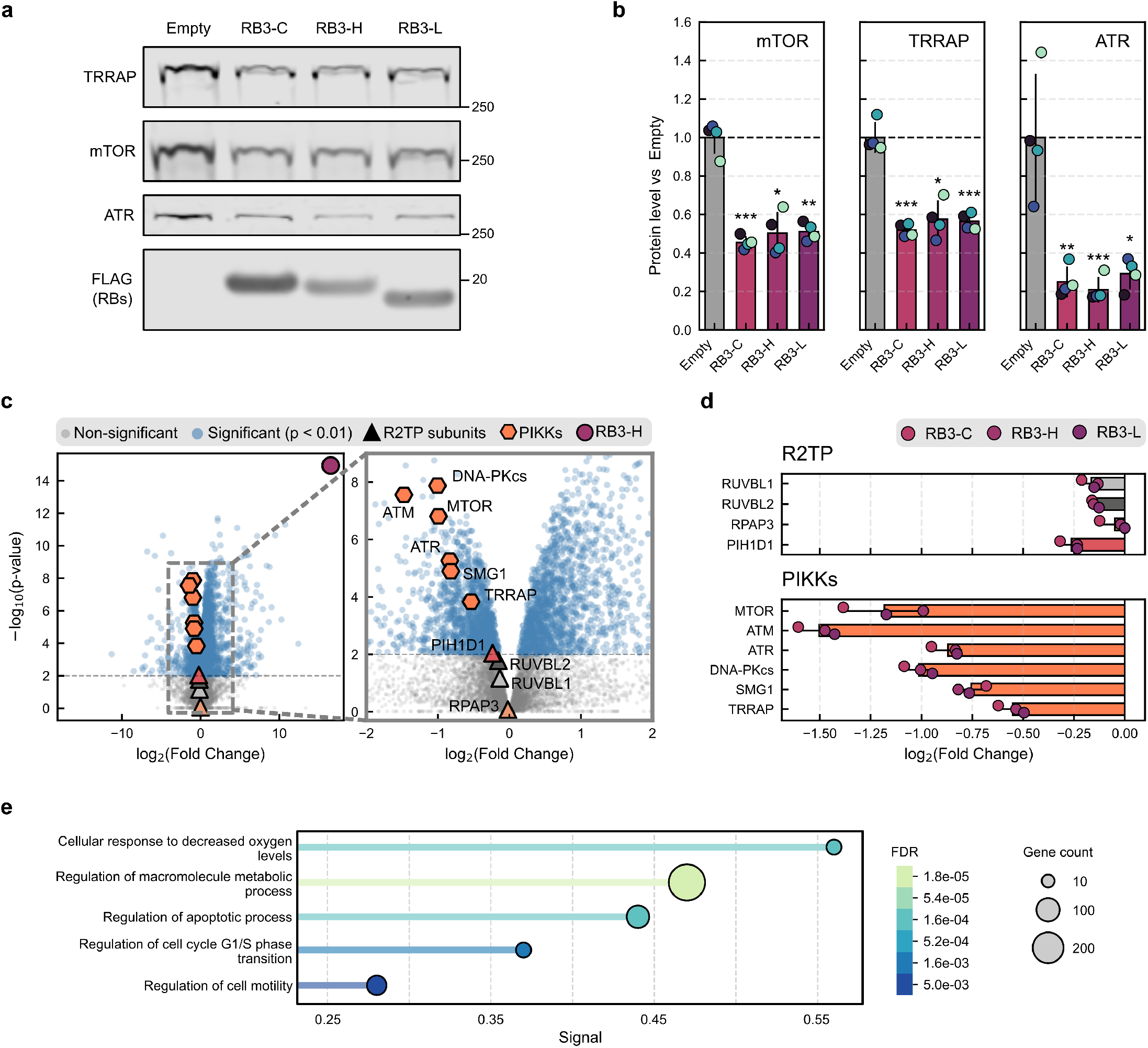
RB3-mediated R2TP disruption reduces cellular abundance of all PIKKs. **a**, Immunoblotting detection of mTOR, ATR and TRRAP after transient expression of FLAG-tagged RB3s, compared to controls transfected with mock vector. **b**, Quantification of panel a, bars represent the mean and vertical lines the SD of the Ponceau-normalized signal expressed relative to the Empty group (dashed line). Overlaid dots represent biological replicates (n=4). The linear axis is shown for interpretability, while significance was calculated on log-transformed values and compared by paired t-tests. *p < 0.05, **p < 0.01, ***p < 0.001. **c**, Volcano plots obtained by global proteomic analysis of HEK293T overexpressing RB3-H relative to cells transfected with mock vector. R2TP subunits and PIKKs are highlighted. Dashed line indicates p = 0.01. p-values were calculated with moderated t-tests (limma) and adjusted by Benjamini–Hochberg (FDR < 0.05). **d**, Relative abundance of R2TP subunits and PIKKs averaged across RB3-C, -H and -L overexpression. Bars indicate average log_2_(FC) and standard deviation. **e**, Functional enrichment analysis of differentially abundant proteins simultaneously identified across RB3-C, -H and -L global proteomics datasets. Genes identified in all three datasets were pooled and classified into pathways by GO-defined biological processes. Differences between RB3 datasets and control conditions were statistically assessed by functional enrichment analysis.

To better characterize the changes in PIKK levels and their consequences, we quantified the differential abundance of the proteomes of HEK293T cells transfected with RB3s compared to mock plasmids by global proteomic analysis. The abundance of R2TP components was not significantly altered irrespective of the RB3 overexpressed, indicating that being part of R2TP is not essential for their stability (**Fig. 5c,d** and **Supplementary Fig. S9a**). Because RPAP3 levels were not significantly altered upon RB3 overexpression, the reduced interaction between RPAP3 and RUVBL1 in the presence of RB3-H (**Fig. 3c**) must be attributed to changes in the amount of assembled R2TP rather than changes in the abundance of its individual components. In contrast, all PIKKs (mTOR, TRRAP, SMG1, DNA-PKcs, ATM, and ATR) showed significantly lower abundance across all three RB3s studied. SMG1 and TRRAP were consistently the least-impacted PIKKs, matching previous reports indicating that SMG1 and TRRAP are the least reliant on R2TP^4^ (**Fig. 5c,d** and **Supplementary Fig. S9a**).

Given the broad regulatory scope of PIKKs, we expected RB-mediated PIKK reduction to result in changes in the abundance of proteins of their respective downstream pathways. We therefore pooled all proteins with consistent differential abundance across all three RB3 datasets and, using STRING-DB^43^, we clustered them to identify physical and functional interactions. Consistent with inhibition of mTORC1, we observed downregulation of anabolism-associated genes like mTORC1 signaling proteins and amino acid transporters, while clusters linked to catabolism, like lysosomal proteins and fatty acid catabolism, appeared upregulated (**Supplementary Fig. S9b**). Also, we observed changes in the abundance of NHEJ, cell-cycle and apoptosis regulation proteins, consistent with altered function of DNA-PKcs, ATR and ATM, essential for a functional DNA damage response (**Supplementary Fig. S9b**,**c**). Moreover, gene set enrichment analyses showed that, indeed, pathways related to cell metabolism and cell cycle were significantly enriched within the pool of differentially abundant proteins identified (**Fig. 5e**).

Together, these data support that the RUVBL1/2-RB interaction, by disrupting R2TP assembly, affects the stability of PIKKs, reducing their cellular abundance and concomitantly affecting their downstream pathways.

## 3. DISCUSSION

Several members of the PIKK family of serine/threonine kinases have become important therapeutic targets: inhibitors of ATR, ATM, and DNA-PKcs sensitize tumor cells to DNA-damaging agents, and mTOR inhibitors are already used in oncology. Despite their therapeutic potential, PIKK inhibitors have faced some limitations, highlighting the interest of alternative strategies to modulate their function^44–49^. One of them is to target their common maturation pathway. R2TP is essential for the stability and maturation of all the members of the PIKK family^3–5^ and therefore, disrupting R2TP assembly could simultaneously impair maturation, and therefore activity, of all PIKKs. Notably, small molecules that inhibit RUVBL1/2 ATPase activity have shown promising effects in cancer models^12–17^. However, since RUVBL1/2 participates in numerous essential cellular functions, including regulation of the Fanconi anemia pathway^19^ and scaffolding of multiple chromatin remodeling complexes^21–23^, ATPase inhibition might perturb many RUVBL1/2-dependent activities rather than selectively disrupting PIKK maturation. This is a problem that extends to other AAA-ATPases that are attractive therapeutic targets, and it has already been proposed that better selectivity could be achieved by specific disruption of protein-protein interactions in certain complexes^24^.

Here, we managed to target specifically the R2TP-related functions of RUVBL1/2 by hampering R2TP assembly, substantially reducing the abundance of all PIKK family members as a result. Our *de novo* designed RBs occupy the RPAP3 binding site in RUVBL1/2, thereby preventing assembly of R2TP and displacing pre-bound RPAP3. The efficiency of RPAP3 displacement correlates with the affinity presented by each RB, showcasing the relevance of design techniques that allow properties of binders to improve iteratively. Our iterative design strategy, relying on partial diffusion and large-scale Protein-MPNN inference^30,31^ to refine backbone geometry and its sequence, respectively, increased their affinity for RUVBL1/2 around 10000×. These results illustrate how computational refinement and small-scale experimental validation, performed iteratively, can optimize binders while substantially reducing the experimental burden of large-scale screening.

Our cryoEM structures showed that backbone adjustments that enable a higher number of contacts between RUVBL1/2 and RBs correlate with affinity increases, as expected. However, the final design stage highlighted a remaining limitation of current protein design approaches. We screened dozens of sequences folding into highly similar backbones (C_*α*_ RMSD between RB3s is < 0.1Å) and highly similar sequences, where most of the residues involved in the RUVBL1/2-RB interaction surface are fully conserved (**Supplementary Fig. S6d**). However, their *K*_*D*_ values span three orders of magnitude. We were unable to find a rationale between amino acid substitution and changes in affinity, highlighting the challenge of predicting the contribution of individual residues, often distal from the interaction surface, on binding properties.

Transient overexpression of RB3s in human cells resulted in up to 70% reduction in the abundance of assembled R2TP, while its individual components remained relatively stable. Consequently, the levels of all PIKKs were significantly reduced. The relative impact on the abundance of each PIKK is consistent with previous reports indicating that maturation of mTOR and ATM are more dependent on the R2TP pathway than SMG1 and TRRAP^4^. In addition, many of the other proteins presenting differential abundance are downstream effectors of PIKK signaling, consistent with impaired PIKK maturation. These results prove that RBs can be used as indirect modulators of PIKK maturation. *De novo* designed proteins are highly stable^34^ and can be genetically encoded, and RBs in particular, feature high affinity and specificity. Given the complex, multi-step and highly interrelated nature of PIKK maturation pathways, RBs represent a valuable tool for their study, enabling the dissection of the R2TP-mediated steps.

One unexpected consequence of designing high-affinity RBs is that they stabilize the ADP-bound conformation in RUVBL1/2, likely precluding the conformational changes required for ATP binding and hydrolysis, as shown by our cryoEM and enzymatic analyses (**Fig. 4**). To our knowledge, this represents the first report of a *de novo* designed allosteric inhibitor acting on a native enzyme. Importantly, this conformational selectivity also provides a mechanistic explanation for the functional specificity of RBs. In non-R2TP RUVBL1/2-containing complexes, such as chromatin remodelers, RUVBL1/2 adopts an ATP-bound-like conformation that remodels the RBD-binding region^27^, rendering it incompatible with RB recognition (**Fig. 4b,c** and **Supplementary Fig. S8**). Furthermore, these complexes engage RUVBL1/2 through its DII-face, opposite to the RBD-binding region^25,50^, so RBs should not compete with them for RUVBL1/2 binding. Together with the lack of nuclear localization signals in RBs, these structural features explain why RBs selectively disrupt R2TP while not affecting chromatin remodeling complexes (**Fig. 4d**). During the preparation of this manuscript, structural work on R2TP-like complexes, R2SP and R2C, showed that they share some architectural features with R2TP, but substitute RPAP3 by other proteins containing an analogous RBD domain capable of binding RUVBL2 through the same region, SPAG1 and CCDC103 (also known as DNAAF19), respectively^26,38,51^. Both complexes participate in the dynein assembly pathway, and mutations in both SPAG1 and CCDC103 have been found in primary ciliary dyskinesia patients due to defective axoneme assembly^52^. Accordingly, given these subunits share RPAP3’s binding site on RUVBL1/2, we observed in our global proteomics data CCDC103 as significantly more abundant (**Supplementary Fig. S9c**) and identified alterations in cell motility and ciliopathy pathways (**Fig. 5e, Supplementary Fig. S9b**), suggesting perturbations in such R2TP-like assemblies. As such RBs also hold promise as tools for further inspecting the participation of R2SP and R2C in the correct assembly of motile primary cilia.

In summary, we designed RBs, *de novo* proteins that bind RUVBL1/2, specifically disrupt R2TP assembly, and reduce the abundance of all PIKKs, with especially pronounced effects on ATM, ATR, and mTOR. We envision RBs as a potential tool for PIKK modulation, with potential clinical and research applications. Inhibition of R2TP and/or RUVBL1/2 activity has recently been explored as a therapeutic strategy for those tumors that are particularly dependent on PIKK activity, as they present a therapeutic window in which their viability can be compromised without major off-target effects. RBs, unlike existing small-molecule inhibitors of RUVBL1/2, affect R2TP-related functions specifically. They also can be genetically encoded, providing flexibility of application to PIKK and R2TP research, and offer room for more rapid and rational engineering, compared with small molecules. The potential of engineered protein binders to disrupt specific interactions is just starting to unfold, and we envision RBs as a useful tool for the R2TP community to begin to exploit it.

## 4. METHODS

### 4.1. Computational design of de novo RUVBL1/2 binders

#### First generation of binders (RB1)

The RUVBL1/2-RBD cryoEM structure 6FO1^25^ was inspected to identify a hydrophobic pocket in the RBD-binding surface to RUVBL2. A single RUVBL1/2 dimer was extracted from the model and trimmed to improve computational efficiency while preserving the RBD-binding interface by removing residues 1–22 and 88–325 of RUVBL1 and residues 1–362 of RUVBL2. The resulting structure was used as input for the BinderFlow standard pipeline, generating batches of eight backbones^30^. For each backbone, a single sequence was designed using ProteinMPNN, followed by backbone energy minimization using the FastRelax protocol^53^. Residues V389, I393, V423, I426, and Y430 were specified as hotspot residues to bias backbone generation towards the RUVBL1/2-RBD interaction surface. Three independent design campaigns were performed, with binder lengths of 60-120, 80-140 and 130-220 residues, and outputs from all campaigns were pooled. Candidate binders were selected by the following criteria: pLDDT_binder > 80, iPAE < 10.

#### Computational refinement of RB1 binders (RB2, RB3)

Experimentally validated binders RB1-A and RB1-B were refined using the BinderFlow Partial Diffusion^35^ pipeline. For each design, its AF2 prediction was used as input to generate batches of eight backbones, sampling similar backbones by applying either 10 or 20 diffusion noise steps and a noise_scale of 1. For each backbone, a single sequence was designed using ProteinMPNN and subjected to FastRelax energy minimization. The top refined binder from each input was selected based on pLDDT_binder>80 and iPAE ranking (RB2-A, iPAE = 8.65 Å; RB2-B, iPAE = 8.0 Å).

The experimentally validated binder RB2-A was further refined using the BinderFlow Sequence Diversity pipeline^30^. The AF2 prediction of RB2-A was used as input, and 10^4^ sequences were generated for the fixed backbone using ProteinMPNN and evaluated using AF2 and PyRosetta. Filter thresholds were iteratively tightened to yield a final set of 27 candidates. Final selection criteria were: iPAE ≤ 7.83 Å; binder_pLDDT ≥ 92; CUTRE < 10^30^; shape_complementarity ≥ 0.55; interface_unsatisfied_hydrogen_bonds ≤ 3; interface_hydrogen_bonds ≥ 4.

### 4.2. Cloning of candidate RUVBL1/2 binders

#### First generation of binders (RB1)

Candidate binder protein sequences were reverse-translated using the reverse translation tool embedded in bfmonitor.py^30^, based on the CodonTransformer library^54^. Overhangs were added in 5’ and 3’ ends for *In vivo* Assembly (IVA) cloning^55^ compatibility and the resulting sequences were ordered as linear DNA fragments (Integrated DNA Technologies). DNA fragments were assembled by IVA cloning into a modified pRSF-Duet plasmid harboring a GFP-10×His C-terminal tag and transformed by heat shock into competent BL21(DE3) *E. coli* for overexpression.

#### Computationally refined binders (RB2, RB3)

Candidate binder protein sequences were reverse-translated and codon-optimized for *E. coli* expression with a custom Python script based on the DNA Chisel library^56^. Adapter sequences were added in 5’ and 3’ ends for compatibility with the Golden Gate Assembly plasmid LM627 (Addgene #191551) as described by Wicky *et al*.^57^. DNA sequences were checked for synthesizability using the DNA manufacturer (Integrated DNA Technologies) web API. Linear DNA fragments were ordered and resuspended in 10 mM Tris-HCl pH 8.0, 0.1 mM EDTA in 96-well plates. 5-*μ*l cloning reactions were performed in 96-well PCR plates for each construct using 1×T4 reaction buffer (New England Biolabs), 20 fmol of LM627, 3 U BsaI-HFv2 (New England Biolabs), 100 U T4 Ligase (New England Biolabs) and 1 ng DNA fragment per well. Cloning reactions were incubated at 37 °C for 20 min followed by 60 °C for 5 min in a thermocycler. Assembled plasmids were transformed into chemically competent BL21(DE3) *E. coli* by adding 1 *μ*l of cloning reaction to 7 *μ*l of cells per well in a 96-well PCR plate on ice, which were sealed with breathable film (Breathe-Easy, Sigma-Aldrich) and incubated for 30 min. Heat shock was performed in a 37 °C water bath for 10 s followed by 2 min on ice. 100 *μ*l of SOC medium were added per well and incubated at 37 °C while shaking at 1000 rpm during 1 hour for heat shock recovery. 900 *μ*l of LB media supplemented with 50 *μ*g/ml kanamycin were dispensed on a 96-deep well block, and cultures were inoculated with 100 *μ*l from the transformed cultures and incubated at 37 °C and 1000 rpm shaking overnight. Glycerol stocks were prepared by mixing 100 *μ*l of saturated cultures in a 1:1 (v:v) ratio with sterile 50% glycerol in LB. The remaining culture volume was used for downstream expression and purification screening.

### 4.3. RUVBL1/2-binder interaction screenings

#### First generation of binders (RB1): pull-down and immunoblotting

Overnight starter cultures were prepared by inoculating 5 ml LB medium supplemented with 50 *μ*g/ml kanamycin (LB-Kan) with glycerol stocks of *E. coli* BL21(DE3) harboring pRSF-Duet–RB1_X–GFP– 10×His plasmids and grown at 37 °C with orbital shaking at 250 rpm. Saturated cultures were diluted 1:1000 into 10 ml LB-Kan and grown at 37 °C with shaking at 250 rpm to mid-log phase (OD_600_ = 0.4-0.6). Protein expression was induced with 0.1 mM IPTG for 3 h at 37 °C. Cells were harvested by centrifugation at 12,000 × *g* for 10 min at 4 °C and pellets were resuspended in 2 ml of lysis buffer (1×PBS, 5% glycerol v:v) supplemented with a cocktail of protease inhibitors (cOmplete EDTA-free, Roche), 2.5 U/ml benzonase (Merck) and lysozyme (0.1 mg/ml) (Sigma-Aldrich). Cells were lysed by sonication on ice for 1 min using 1-s pulses at 38% amplitude (Q125 Sonicator, QSonica), and lysates were clarified by centrifugation at 12,000 × g for 30 min at 4 °C. Clarified supernatants were incubated with 50 *μ*L Ni^2+^–NTA agarose resin (ABT) pre-equilibrated in wash buffer (1× PBS, 50 mM imidazole), followed by two washes with 10 column volumes (CV) of wash buffer. Purified RUVBL1/2 complex (100 *μ*l at 0.05 mg/ml), prepared as described in Lopez-Perrote *et al*.^37^, was added to the resin-bound RB1 candidate binders and incubated on ice for 30 min. The resin was washed twice with 10 CV of wash buffer, and bound proteins were eluted with 50 *μ*l wash buffer supplemented with 500 mM imidazole. Samples were resolved by SDS-PAGE and transferred to nitrocellulose membranes in transfer buffer (50 mM Tris, 380 mM Glycine, 20% ethanol) for 2 h at 300 mA and 4 °C. Membranes were blocked by incubation with 3% BSA in WB buffer (20 mM Tris pH 7.5, 150 mM NaCl, and 0.05% Tween-20), washed extensively with WB buffer and probed with anti-RUVBL2 monoclonal antibody (OriGene #CF504284) according to the manufacturer’s indications.

#### Computationally refined binders (RB3): high-throughput pull-down

— 400 *μ*l saturated cultures grown overnight after Golden Gate Assembly were used to inoculate 3.6 ml of Terrific Broth supplemented with 50 *μ*g/ml kanamycin per well in 24-ml deep well blocks, totaling 4 ml of culture per design. Cultures were incubated at 37 °C with shaking at 275 rpm for 2 h. Protein expression was induced by addition of IPTG to a final concentration of 1 mM. Following induction, cultures were incubated for an additional 4 h at 37 °C with shaking at 275 rpm. Cells were harvested by centrifugation at 4000 × *g* for 10 min. Bacterial pellets were resuspended in 400 *μ*l of Bacterial Protein Extraction Reagent (B-PER, ThermoFisher), supplemented with 0.1 mg/ml lysozyme, 50 U/ml benzonase and protease inhibitors (cOmplete EDTA-free, Roche), vortexed until homogeneous, and incubated at 18 °C with shaking for 15 min. Lysates were clarified by centrifugation at 3200 × *g* for 60 min at 4 °C. Fritted 96-well plates were loaded with 100 *μ*l regenerated Ni^2+^-NTA resin (ABT) per well on a plate vacuum manifold. The resin was equilibrated by drawing four volumes of 200 *μ*l wash buffer (20 mM Tris-HCl pH 8.0, 300 mM NaCl, 25 mM imidazole) through each well. Clarified lysates were applied to the resin in each well and allowed to flow through the resin by gravity. The resin was washed four times with 150 *μ*l wash buffer per well under a gentle vacuum. Residual buffer was removed by blotting the plate on absorbent paper. For expression screening, proteins were immediately eluted in 100 *μ*l elution buffer (20 mM Tris-HCl pH 8.0, 300 mM NaCl, 500 mM imidazole) onto a V-bottom 96-well deep well block and analyzed by SDS-PAGE. Large scale protein production for downstream biophysical analysis was performed following the same protocol, but scaling all reagents 3-fold, for a total volume of *E. coli* cultures of 12 ml.

For interaction screening, purified, untagged RUVBL1/2 complexes^37^ were diluted in wash buffer to 5 *μ*M. 95 *μ*l of RUVBL1/2 were added to each well. The plate was sealed and incubated at 4 °C for approximately 25 min to allow complex formation. Following incubation, the plate was placed back onto the manifold and washed five times with 150 *μ*l wash buffer. Bound proteins were eluted twice by drawing 100 *μ*l elution buffer into a V-bottom 96-well deep-well block. Eluted proteins were analyzed by SDS-PAGE. A negative control consisting of resin exposed only to RUVBL1/2 protein without lysate was included.

### 4.4. Size exclusion chromatography (SEC)

A Superose 6 Increase 3.2/300 column was equilibrated in 1× PBS. Protein complexes consisting of RUVBL1/2 and individual binders were prepared at a 1:5 molar ratio (RUVBL1/2 dimer:binder) and incubated for 40 min at 25 °C prior to injection. All runs were performed at a constant flow rate of 0.06 ml/min monitoring UV absorbance at 280 nm and collecting 50 *μ*l fractions. Binder-only and RUVBL1/2-only control samples were prepared and injected under identical conditions.

### 4.5. Isothermal titration calorimetry (ITC)

Purified RUVBL1/2 and binder samples were simultaneously buffer-exchanged into 1× PBS using 3.5 kDa molecular weight cutoff dialysis cassettes (Slide-A-Lyzer, ThermoFisher) overnight at 4 °C to ensure identical buffer composition. Following dialysis, samples were recovered and protein concentrations were determined by UV absorbance. ITC was performed at 30 °C using a MicroCal iTC200 (Malvern Instruments, UK). Binders in concentrations ranging 120 to 220 *μ*M were titrated into RUVBL1/2 solution at 15 *μ*M by 19 × 2-*μ*l injections spaced by 150 s delays and continuous stirring at 1000 rpm. When possible, blank contribution was estimated from the five last injections and subtracted from the protein-protein titration data. Data were analyzed using Origin 7.0 (OriginLab) or PEAQ-ITC (Malvern Panalytical) using a one-site binding model.

### 4.6. Biolayer interferometry (BLI)

Untagged, purified RUVBL1/2 complexes were biotinylated using the ChromaLink Biotin Protein Labeling Kit (VectorLabs, #B-9007105K) according to the manufacturer’s indications. The BLI screening assays were performed on an 8-channel Octet RED96 system (ForteBio) at 30 °C, the finer determination of RB3-C, -H and -L kinetic parameters was performed on an 8-channel Octet R8 system (Sartorius). Biotinylated RUVBL1/2 complexes were immobilized onto Streptavidin (SA)-conjugated BLI tips (SA biosensors, Sartorius) for 30 s at a 30 nM concentration. SA tips were dipped into 1× PBS for 180 s for removal of excess RUVBL1/2 complexes and establishing a sensorgram baseline. The association phase was performed in dilutions of purified binders in 1× PBS at the concentrations indicated in the figure for 240 s. Dissociation was recorded for 600 s in 1× PBS. Baseline signal was averaged and removed from the whole sensorgram trace. Kinetic and equilibrium constants were globally fitted across all concentrations to a 1:1 binding model using Octet’s analysis software (Sartorius). All experimental traces shown were subjected to Savitzky-Golay filtering for noise averaging. Sensorgrams with low signal-to-noise ratio were removed from analysis.

### 4.7. CryoEM of RUVBL1/2-binder complexes

#### Sample preparation and data collection

20 *μ*M RUVBL1/2 were incubated with 120 *μ*M of either RB1-A, RB2-A, RB3-C or RB3-H, or 180 *μ*M of RB3-L, during dialysis into 1× PBS using 3.5 kDa molecular weight cutoff dialysis cassettes (Slide-A-Lyzer, ThermoFisher) for 5 hours at 4 °C. Quantifoil R1.2/1.3 Cu 300 or Quantifoil R1/1 Cu 300 were glow discharged at 15 mA 0.1 bar for 45 s (GlowQube, Quorum Technologies) with residual atmospheric air. 3 *μ*l of sample were applied, blotted for 3 to 4 s and vitrified in a liquid ethane-propane mixture using an FEI Vitrobot Mark IV (Thermo Scientific) operating at 90% humidity and 4 °C.

Data for the RUVBL1/2-RB2-A complex were collected on a 300 kV Titan Krios G4 (Thermo Scientific) electron microscope located at the Basque Resource for Electron Microscopy (BREM, Spain) equipped with a K3 detector (Gatan) operated in counting mode and a Bio-Continuum Imaging Filter with 20 eV slit width controlled via EPU (Thermo Scientific), at nominal magnification of 81,000× (1.05 Å/px), a total dose of 52.5 e^-^/Å^2^ fractionated in 50 frames, and nominal focus ranging from -0.7 to -1.7 *μ*m. Data for the RUVBL1/2-RB1-A, -RB3-C, -RB3-H and -RB3-L complexes were collected on a Titan Krios G3 (Thermo Scientific) located at the UK’s national Electron Bio-Imaging Centre (eBIC), equipped with a Falcon IV direct detector and a SelectrisX energy filter with 10 eV slit width controlled via EPU, a 50 *μ*m C2 aperture at nominal magnification of 130,000× (0.931 Å/px) — except for the RB3-C dataset, collected at 165,000× (0.729 Å/px) — a total dose of 40.1 e^-^/Å^2^ fractionated in 46 frames and nominal focus ranging from -0.8 to -2.0 *μ*m.

#### CryoEM data processing

Each dataset was processed independently, following the pipeline shown in **Supplementary Fig. S4 and S5**, where number of movies and particles per step are indicated. Raw movies were motion-corrected, dose-weighted and aligned using the MotionCor2^58^ algorithm implementation in RELION-5.0^59,60^. CTF parameters of motion-corrected micrographs were estimated with CTFFIND4.1^61^. Micrographs were curated using relion_analyse.py^62^ to remove low quality or ice-contaminated images and transferred to cryoSPARC v4.7^63^. Particles were picked repeatedly with different algorithms (blob picking, template picking and TOPAZ^64^) attempting to identify underrepresented particle orientations. Duplicate particles were removed according to their picking coordinates and were iteratively curated by 2D classification and averaging. Particles belonging to 2D classes that featured secondary structure elements were used for multiclass *ab initio* reconstruction in cryoSPARC v4.7. A volume resembling the expected RUVBL1/2 hexamer bound to three helical densities in the RBD-binding surface and a noisy volume resulting from misalignment during *ab initio* reconstruction was used for supervised 3D classification in cryoSPARC (Heterogeneous refinement job) or RELION-5.0 (3DClass job). Particles yielding volumes consistent with RUVBL1/2 complexes were pooled and used for high-resolution gold-standard angular refinement (Homogeneous refinement job). 3D volumes were inspected for local occupancy using OccuPy^39^. Aligned particles were subjected to a round of unsupervised 3D classification without alignment (3D Class job in cryoSPARC, 3DClass job turning off alignments in RELION) and refined again, yielding high-resolution volumes in all cases. While the nominal resolution of the maps remained similar, optical aberration, per-particle CTF, defocus^65–67^ and motion^68^ corrections improved notably the interpretability of the map, particularly in the binder region. To further improve map quality, we aligned particles to the C3 symmetry axis and performed symmetry expansion, and 3D classification without alignment in cryoSPARC or RELION to identify the highest-quality asymmetric subunits. Several rounds of 3D classification and 3D local refinement were performed until no heterogeneity was detected, yielding the final volumes.

### 4.8. Model building

Raw cryoEM refined maps were postprocessed using DeepEMhancer^69^ only to facilitate side chain visualization. PDB 6FO1 and AF2 predictions for each binder were rigid-body fitted into the cryoEM densities using ChimeraX^70^. Side-chain positions were manually adjusted and loops in the DII domain of RUVBL1 were built *de novo* in ISOLDE^71^. All models were iteratively real-space refined using ServalCat^72^ within CCP-EM Doppio’s interface. Models were validated using MolProbity^73^.

### 4.9. RUVBL1/2-RBD competition assay

Preassembled RUVBL1/2–RBD complex, carrying a C-terminal Strep-Tag in RBD, was immobilized on 20 *μ*l of StrepTactin XT resin pre-equilibrated in PBS for 1 h at 4 °C under rotation. Samples were transferred to spin columns and unbound RUVBL1/2–RBD was removed by 35 resin volumes washing with 1×PBS. RB was freshly diluted from concentrated stocks to final concentrations of 0, 5, 10, 30 and 60 *μ*M. Resin-bound RUVBL1/2–RBD was incubated with 40 *μ*l of the respective binder solution for 1 h at 4 °C with shaking, and competition wash was collected. RB-containing washes were analyzed by SDS-PAGE and visualized by Coomassie staining. Competition was assessed as a RB-concentration-dependent increase of RUVBL1/2 in the RB-containing washes.

### 4.10. Cell culture and transfection

Each binder’s ORF was subcloned from their respective bacterial expression plasmid into a pRK5 vector containing a C-terminal 3xFLAG tag for overexpression in human cells using IVA cloning^55^. HEK293T cells were cultured in Dulbecco’s Modified Eagle Medium (DMEM) supplemented with 10% fetal bovine serum at 37 °C and 5% CO_2_. 6 × 10^6^ cells were transiently transfected using Lipofectamine 2000 (Invitrogen) and 15 *μ*g of each plasmid according to the manufacturer’s indications. 24 h after transfection, cells were washed. 72 h after transfection cells were harvested by resuspension in 1× PBS and centrifugation at 800 × g for 5 min. Pellets were either flash-frozen and stored at -80 °C or processed immediately. All experiments were carried out using cells confirmed to be free of mycoplasma contamination, monthly tested using qPCR by the CNIO’s Monoclonal Antibodies Unit.

### 4.11. PIKK immunoblotting analysis

Cells were harvested and lysed at 4 °C on a shaker for 20 min using Urea buffer (50 mM Tris-HCl pH 7.5, 8 M urea, and 1% CHAPS) with an incubation step of 30 min at 4 °C. After lysis, the protein fraction was isolated by centrifugation at 13,200 rpm for 10 min at 4 °C and protein concentrations were determined by Bradford Protein Assay (Bio-Rad). Protein extracts were mixed with NuPAGE LDS (LifeTechnologies) supplemented with 10 mM dithiothreitol (DTT, Sigma) and were incubated at 99 °C for 10 min. Samples were resolved on precast 4-20% gradient SDS-polyacrylamide gels (Invitrogen) and transferred to nitrocellulose membranes in transfer buffer at 100 V for 1 h at 4 °C. Membranes were stained with Ponceau S, destained and blocked with 5% non-fat dry milk dissolved in TBS-T. Membranes were probed with primary antibodies against mTOR (Cell Signaling #2983, 1:1000 dilution), ATR (Serotec, #AHP384, 1:500 dilution), TRRAP (Cell Signaling, #3967, 1:1000 dilution), or FLAG (Sigma, #F1804, 1:1000 dilution). Fluorophore-conjugated secondary antibodies were used for detection, using the LI-COR Odyssey CLx imaging system (LI-COR Biosciences).

### 4.12. Co-immunoprecipitation

Each cell pellet was resuspended in lysis buffer (50 mM HEPES pH 7.5, 150 mM NaCl, 1% Triton X-100, 1 mM EDTA, 5% glycerol) supplemented with protease inhibitor cocktail (cOmplete EDTA-free, Roche) and 2 U/ml benzonase. Lysates were incubated for 50 min at 4 °C with rotation and clarified by centrifugation at 16,000 × *g* for 45 min at 4 °C. Total protein concentrations were determined by Bicinchoninic Acid Assay (BCA) assay. For RUVBL1 co-immunoprecipitation, Protein G Dynabeads (Invitrogen) were incubated for 35 min at 25 °C with rotation with polyclonal anti-RUVBL1 antibody (Proteintech #10210-2-AP), washed and equilibrated in lysis buffer. 300 *μ*g of protein from whole cell lysates were incubated with antibody-coupled beads for 1 h at 4 °C. For RB3-H co-immunoprecipitation, anti-FLAG M2 magnetic beads (Millipore) were equilibrated in lysis buffer. 300 *μ*g of protein from whole cell lysates were incubated with beads overnight at 4 °C with rotation. Beads were washed six times with six resin volumes in wash buffer (50 mM HEPES pH 7.5, 150 mM NaCl), and resin-bound proteins were flash-frozen in liquid N_2_ for downstream processing.

### 4.13. ATP hydrolysis assays

ATPase activity of RUVBL1/2 in presence and absence of interaction partners was measured in a continuous spectrophotometric pyruvate kinase-lactate dehydrogenase-coupled enzymatic assay, which couples the regeneration of the hydrolyzed ATP to the oxidation of NADH^74^. Abs_340_ from NADH was measured using a CLARIOstar Plus plate reader (BMG LabTech), and the slope of the curves was used to estimate the ATP hydrolysis rates. Assays were performed at 37 °C in 100 *μ*l reactions in reaction buffer (50 mM Tris-HCl pH 7.4, 150 mM NaCl, 20 mM MgCl_2_) containing 2 mM phosphoenolpyruvate, 0.5 mM NADH, 0.04 U/*μ*l pyruvate kinase, 0.05 U/*μ*l lactate dehydrogenase (Sigma-Aldrich), 5 mM ATP and 2.5 *μ*M RUVBL1/2 dimers. RUVBL1/2 interaction partners were used at 15 *μ*M for binders and RBD (1:6 molar ratio) or 5 *μ*M for full-length RPAP3 (1:2 molar ratio). The reactions were monitored for 60 min. ATP turnover was calculated for time intervals in which Abs_340_ decreased linearly (time points 10 to 40 min) and fitted to a linear model by least squares approximation. Slopes were normalized to the average slope of buffer in absence of RUVBL1/2. All assays were performed at least in triplicate.

### 4.14. Mass spectrometry analyses

#### Protein extraction and digestion

For global proteomics analysis, cell cultures were maintained, transfected and harvested as described above. Pellets were lysed in buffer containing 4% SDS and 50 mM Tris-HCl pH 8.0 and incubated at 95 °C for 15 minutes with agitation (1200 rpm). Lysates were clarified by centrifugation to remove insoluble material, and protein concentration was determined by BCA. For co-immunoprecipitation analysis, resin-bound proteins were eluted in the presence of 2% SDS and at room temperature for 15 min. Proteins were reduced with tris(2-carboxyethyl)phosphine (TCEP, 10 mM) at 37 °C for 30 min and alkylated with 30 mM chloroacetamide at 37 °C for 30 min. 25 *μ*g of total protein per sample were processed using an automated protein clean-up and digestion workflow implemented on a KingFisher Apex system with MagReSyn® Hydroxyl magnetic beads (ReSyn Biosciences), using a protein-to-bead ratio of 1:2. Proteins were digested overnight (16 h) at 37 °C in 50 mM HEPES pH 8.25 supplemented with 5 mM CaCl_2_, using trypsin and Lys-C proteases at enzyme-to-protein ratios of 1:50 and 1:100, respectively. Digestion was quenched by acidification with trifluoroacetic acid to a final concentration of 1% (v/v).

#### Peptide cleanup and LC–MS/MS acquisition

500 ng of peptides per sample were desalted and concentrated using Evotip® Pure tips (Evosep). Samples were analyzed using an Evosep One LC system operated with a pre-programmed WisperZoom gradient (20 samples per day) and coupled online to an Orbitrap Astral mass spectrometer (Thermo Fisher Scientific) via an EASY-Spray™ ion source. Peptides were separated on a 15-cm Aurora Elite column (75 *μ*m inner diameter, 1.7 *μ*m particles). The spray voltage was set to 1.6 kV, funnel RF level to 40, and capillary temperature to 280 °C. Data-independent acquisition was performed with full MS scans acquired in the Orbitrap analyzer at a resolution of 240,000 over an m/z range of 380–980, using an AGC target of 500% and a fixed cycle time of 0.6 s. Fragment ion spectra were acquired in the Astral analyzer at a resolution of 80,000 with a maximum injection time of 3 ms and an AGC target of 100%. A total of 300 isolation windows of 2 Th were used across the 380–980 m/z range. Fragmentation was performed by higher-energy collisional dissociation with a normalized collision energy of 27%.

#### Statistical analysis

Downstream data analysis was performed in R (v4.3.1). Protein intensity values were log_2_-transformed prior to analysis, and proteins were retained if at least 70% of samples contained measured values in at least one condition of the comparison. Normalization and missing-value handling were performed using the DAPAR (v1.34.6) framework. Partially observed values (POV), defined as proteins with at least one valid measurement within a condition, were imputed using the Structured Least Squares Algorithm (SLSA) with selec = 5000 and otherwise default parameters. Proteins missing entirely within a condition were imputed using a quantile-based approach implemented in the imp4p (v1.2) package. LOESS normalization was applied across samples within each pairwise comparison.

Differential protein abundance analysis was performed using the limma package. For each pairwise comparison, statistical significance was assessed using moderated t-tests, and p-values were adjusted for multiple testing using the Benjamini–Hochberg method. Proteins were considered differentially abundant at a false discovery rate threshold of < 5%. Fold-change cutoffs were applied in a comparison-dependent manner, selected to reflect the distribution of effect sizes and statistical confidence in each comparison. To avoid significance driven predominantly by imputed values, p-values were set to 1 (non-significant) for proteins in which fewer than 70% of samples in the condition with increased abundance contained valid (non-imputed) measurements. Data are available via ProteomeXchange with identifier PXDXXXX.

### 4.15. Clustering and enrichment analysis

Proteins showing significant differential abundance across all three global proteomics datasets (RB3-C, -H and -L) were pooled and submitted as gene symbols to STRING^43^. Interactions were retrieved using the full STRING network and a minimum required interaction score of 0.4. The resulting network was partitioned into modules with the MCL algorithm as implemented in STRING. Ten clusters with ≥4 members and interpretable, non-redundant annotation were selected for display in the figures. Functional enrichment of the consensus set was performed with the enrichment module built into STRING, which tests each annotation term for over-representation among the submitted proteins against the whole human genome, correcting for multiple testing by Benjamini–Hochberg. Terms were ranked by STRING’s *signal* score, combining enrichment and statistical significance. Terms were grouped by similarity at a threshold of 0.7 to remove redundancy inherent to GO hierarchy.

## 5. ACKNOWLEDGEMENTS

We acknowledge the assistance of Dr. Marta Isasa and Dr. Fernando García from the CNIO Proteomics facility for proteomics data collection and analysis, and Dr. Clara Santiveri and Dr. Ramón Campos from the CNIO Spectroscopy and NMR Unit for assistance with ITC data collection and analysis. We also acknowledge the assistance of Dr. José Antonio Infantes from the ISCIII during BLI data collection. CryoEM data were obtained at the Diamond Light Source cryoEM facility at the UK’s National Electron Bio-Imaging Centre (eBIC) under BAG “Stop cancer – structural studies of macromolecular complexes involved in cancer by cryoEM” (BI39322) and at the Basque Resource for Electron Microscopy located at Instituto Biofisika (UPV/EHU, CSIC), supported by the Department of Science, Universities and Innovation and the Innovation Fund of the Basque Government, with additional support from MCIN (Recovery,

Transformation and Resilience Plan) and the Basque Government “Biotechnology Complementary Plan Applied to Health” with funding from European Union NextGenerationEU [PRTR-C17.I1, PRTR-C17.I01.P01.S13, AAAA_ACG_AY_2539/22_05]. This work benefited from access to EMBL Hamburg for BLI measurements, part of the Instruct-ERIC research infrastructure through a canSERV project to R.F.-L. (PID: 38599). Financial support was provided by the European Union’s Horizon Europe research and innovation programme under grant agreement number 101058620 canSERV.

This work was funded by grants supported by the State Research Agency (AEI, 10.13039/501100011033) from the Spanish Ministry of Science and Innovation (PID2023-146110NB-I00 to OL and CNS2023-143762 to RFL), cofunded by ERDF. O.L. was also supported by the program of R&D activities TecNanoBio-CM (TEC-2024/TEC-158) granted by the Autonomous Region of Madrid through the “Dirección General de Investigación e Innovación Tecnológica”. This work was developed at the CNIO funded by the Health Institute Carlos III (ISCIII) and the Spanish Ministry of Science and Innovation. N.G.-R. was supported by a Boehringer Ingelheim PhD fellowship and a Fulbright PhD Research Fellowship.

## Funding of open access

This will be covered by grants supported by the State Research Agency (AEI, 10.13039/501100011033) from the Spanish Ministry of Science and Innovation (PID2023-146110NB-I00 to O.L. and CNS2023-143762 to R.F.-L.), split in half for each of the grants.

## 6. AUTHOR CONTRIBUTIONS

**N.G.-R**.: Conceptualization, Data Curation, Investigation, Formal Analysis, Software, Visualization, Writing – original draft, Writing – review & editing. **A.L.P**.: Investigation, Formal Analysis. **C.R**., **J.A-L**., **M.F**., **E.B**., **E.C**., **I.H**., **G. L-P**., **M. M**.: Investigation. **C.C**.: Software. **O. F-C**.: Funding Acquisition, Supervision. **O.L**., **R.F.-L**.: Conceptualization, Funding Acquisition, Project Administration, Supervision, Writing – original draft, Writing – review & editing.

## 8. DATA AND MATERIALS AVAILABILITY

Materials are available upon request to O.L. and R. F.-L., subject to standard material transfer agreements. CryoEM maps and atomic models have been deposited in the Electron Microscopy Data Bank (EMDB) and Protein Data Bank (PDB) under the following accession codes: RUVBL1/2–RB1-A, EMD-XXXX and XXXX; RUVBL1/2– RB2-A, EMD-XXXX and XXXX; RUVBL1/2–RB3-C, EMD-XXXX and XXXX; RUVBL1/2–RB3-H, EMD-XXXX and XXXX; and RUVBL1/2–RB3-L, EMD-XXXX and XXXX. The mass spectrometry proteomics data have been deposited to the ProteomeXchange Consortium via the PRIDE partner repository with the dataset identifier PXDXXXX.

