## Supplementary Material for "Disruption of R2TP-mediated PIKK maturation with *de novo* designed binders"

\*These authors contributed equally.

Supplementary figures and table for the accompanying manuscript. Typeset with the rho-class L<sup>A</sup>T<sub>E</sub>X template.

### 1. SUPPLEMENTARY FIGURES

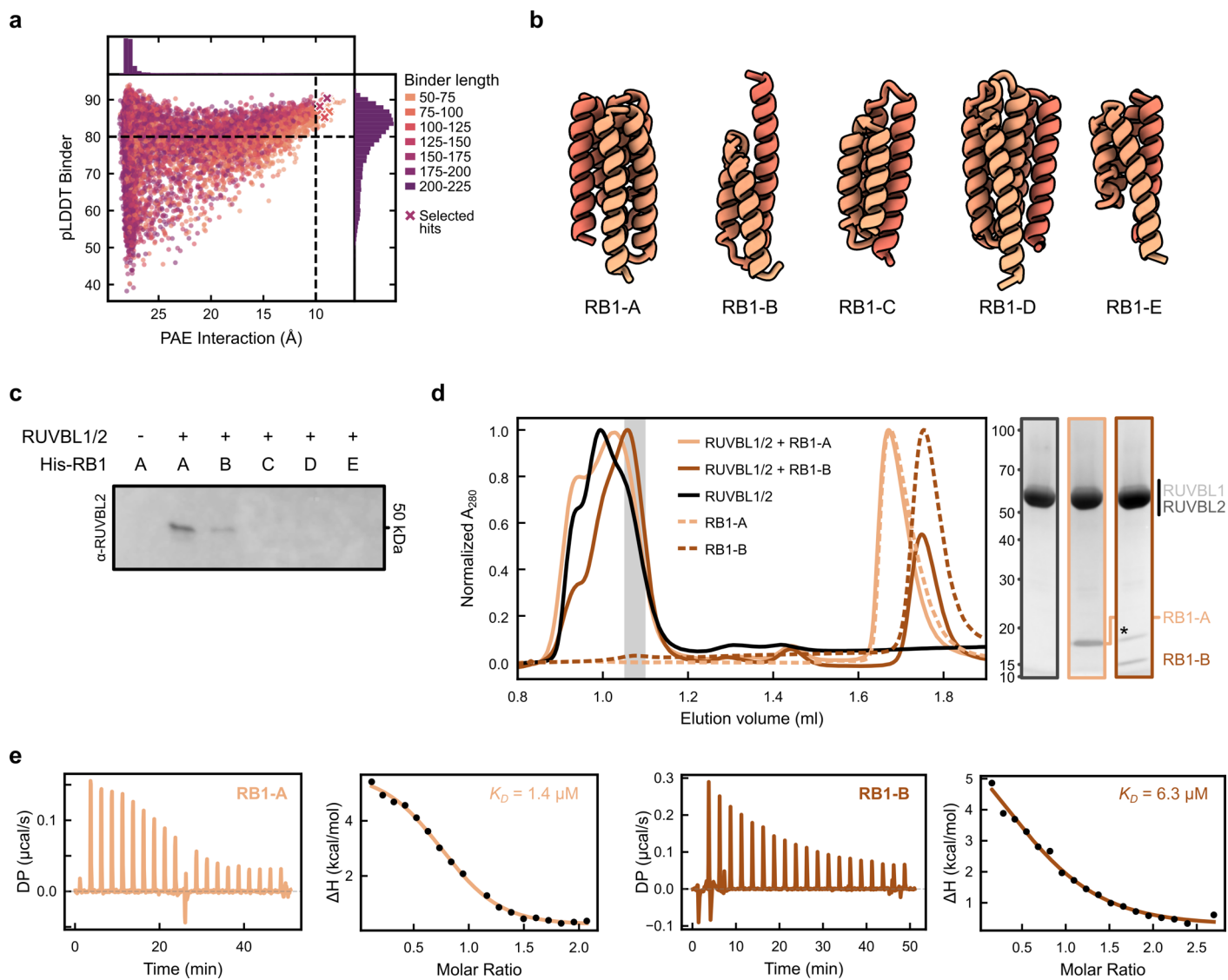

**Supplementary Figure S1. Initial RB design campaign.** **a**, Scatter plot representing each of the candidates generated during the first round of computational design. The dashed black lines represent filtering thresholds: iPAE ($< 10 \text{ \AA}$) and pLDDT\_binder ($> 80$). Designs selected for experimental validation are highlighted as crosses. **b**, Designs selected for experimental validation shown as cartoons. **c**, Immunoblot detection of RUVBL2 in eluates from pull-downs of purified RUVBL1/2 complexes incubated with resin-bound His-tagged RB1 designs. **d**, SEC profiles of RUVBL1/2 purified complexes after incubation with RB1-A and -B designs. The shadowed area indicates the elution volumes contained in the SDS-PAGE fractions shown on the right. Asterisk (\*) indicates a contaminant from the RB1-B purification. RUVBL1 and RUVBL2 have similar electrophoretic mobility, so their bands are not easily resolved in SDS-PAGE. **e**, ITC thermograms and integrated heats (shown side by side for each case) of the interaction between RUVBL1/2 and RB1-A (left) and RB1-B (right).

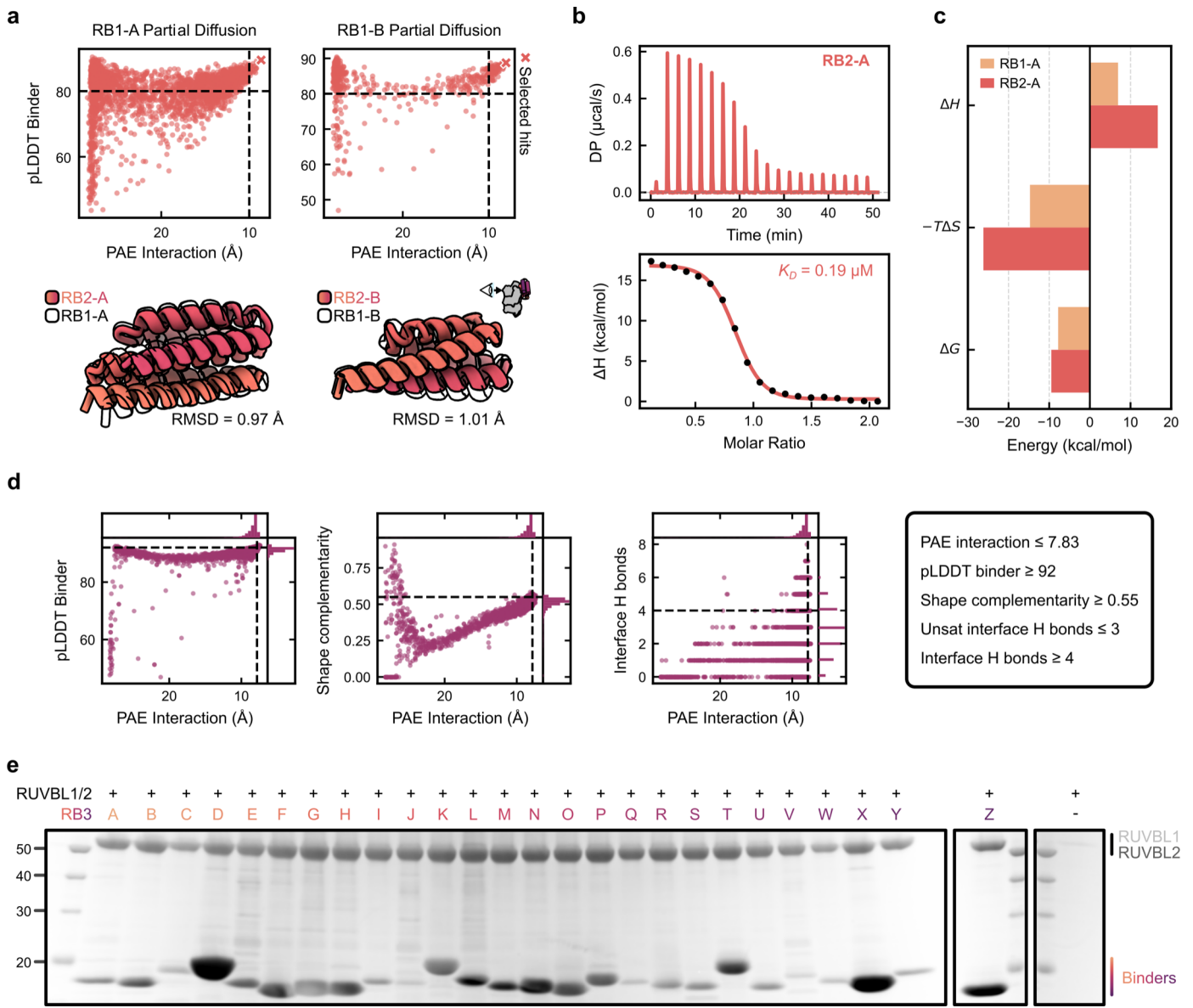

**Supplementary Figure S2. RB refinement campaigns.** **a**, Scatter plots representing the candidates generated during the second round of design, using partial diffusion (top), and cartoon representation of aligned RB1s and their respective RB2s (bottom). The two campaigns are displayed, using RB1-A (left) and RB1-B (right) as starting points. The dashed black lines represent the iPAE ($< 10 \text{ \AA}$) and pLDDT\_binder ($> 80$) thresholds used for filtering. The designs selected for experimental validation are highlighted in the scatter plots as crosses. The RMSD values displayed were calculated between the backbones' $C_\alpha$ atoms. **b**, ITC thermogram (top) and integrated heats (bottom) of the RUVBL1/2-RB2-A interaction. **c**, Comparison of the thermodynamic profiles of RB1-A and RB2-A, with the data obtained from Supplementary Figure S1e and panel b from this figure. **d**, Scatter plots representing the candidates generated during the third round of design, Sequence Diversity RB2 refinement. pLDDT\_binder, shape complementarity and number of interface H bonds are plotted against iPAE. The black, dashed lines represent the threshold values for hit selection also indicated in the grey text box. **e**, SDS-PAGE analysis of the eluates from RUVBL1/2-RB3 pull-down screening. Purified RUVBL1/2 was flown through Ni-NTA resin previously bound to His-tagged RB3s. The presence of RUVBL1/2 bands indicates interaction with the respective RB3s.

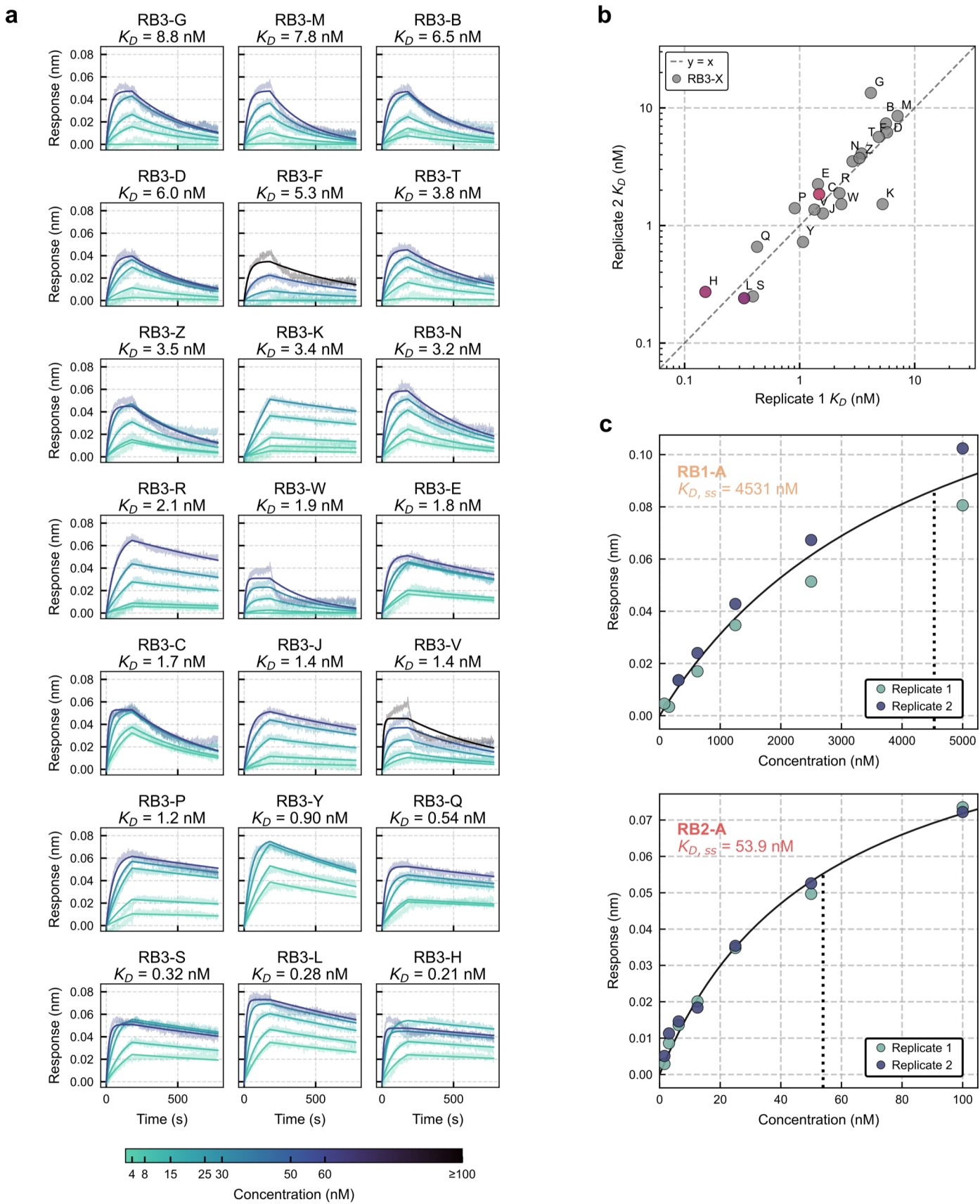

**Supplementary Figure S3. Affinity characterization of RUVBL1/2-RB3 interactions.** **a**, BLI traces of all RB3 candidates binding to biotinylated RUVBL1/2 immobilized onto streptavidin-conjugated sensors. Solid lines represent kinetic fittings; transparent lines indicate experimental traces ($n=2$). All traces were smoothed using the Savitzky-Golay algorithm. **b**, Scatter plot of correlation of $K_D$ values determined by global fitting of independent replicates. The dashed line, $x=y$, indicates perfect correlation. **c**, BLI steady-state analysis of RB1-A (top) and RB2-A (bottom). Black lines indicate fitting to the binding isotherm.

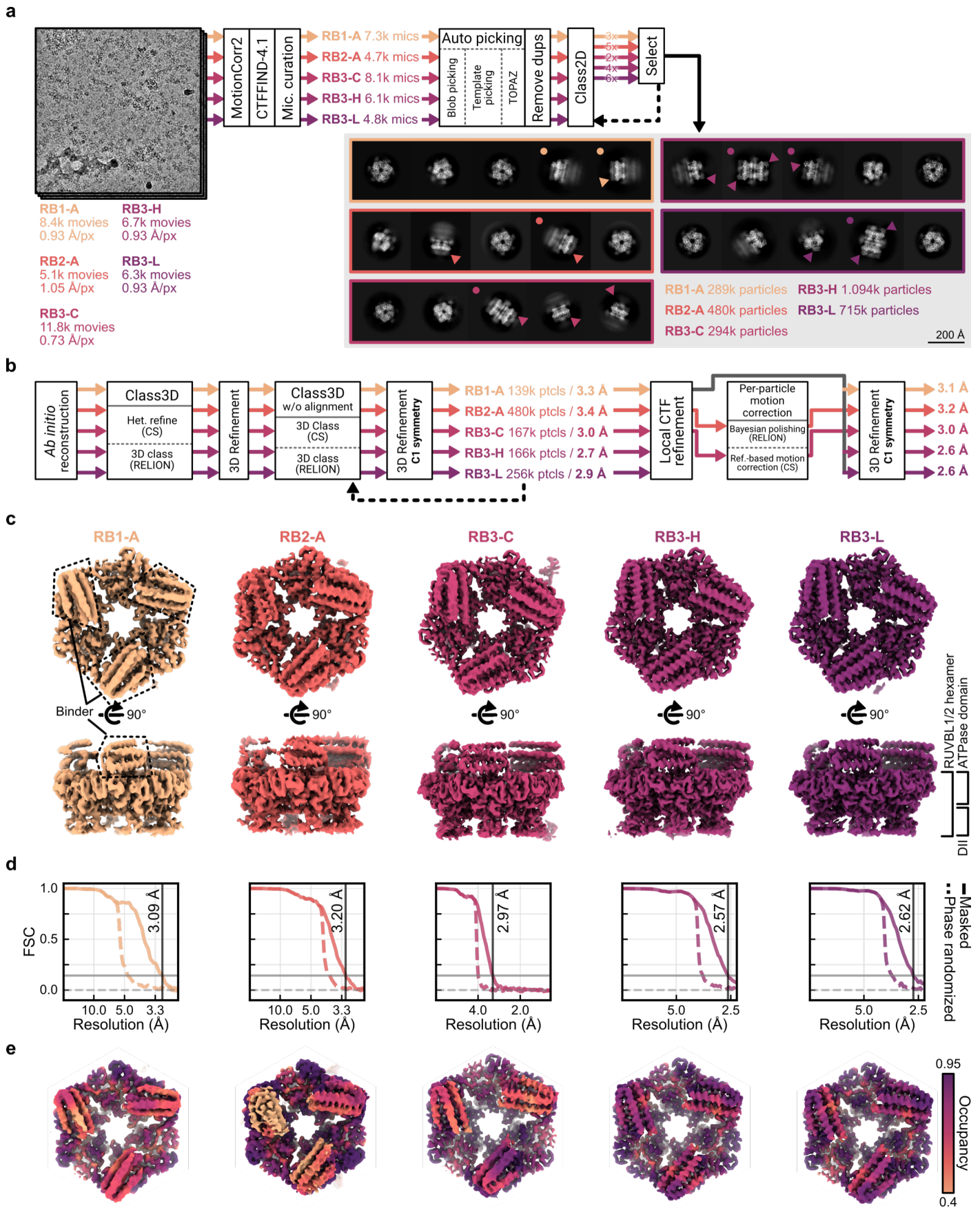

**Supplementary Figure S4. CryoEM of RUVBL1/2-RB complexes.** **a**, Data collection and preprocessing pipeline followed for each of the RB datasets. 2D averages highlighted with dots indicate dodecameric RUVBL1/2 side views. Triangles overlaid on the 2D averages point to the RB helical bundle bound to the ATPase face of RUVBL1/2. **b**, 3D reconstruction pipeline followed for each RUVBL1/2-RB complex. **c**, Density surface representation of all raw, unsharpened RUVBL1/2-RB complexes. The dashed lines in the leftmost column indicate the densities attributed to RB α-helical bundles. **d**, Gold-standard FSC plots of each of the final 3D reconstructions. X-axes are plotted in reciprocal scale ($1/x$), but x-axis labels correspond to real-space frequency values. Horizontal grey lines indicate FSC = 0 (dashed) and FSC = 0.143 (solid). Vertical black line indicates the resolution at FSC = 0.143. **e**, Density surface representation of each RUVBL1/2-RB complex colored by local scale, as calculated by OccuPy. All panels are color-coded according to each RUVBL1/2-RB dataset.

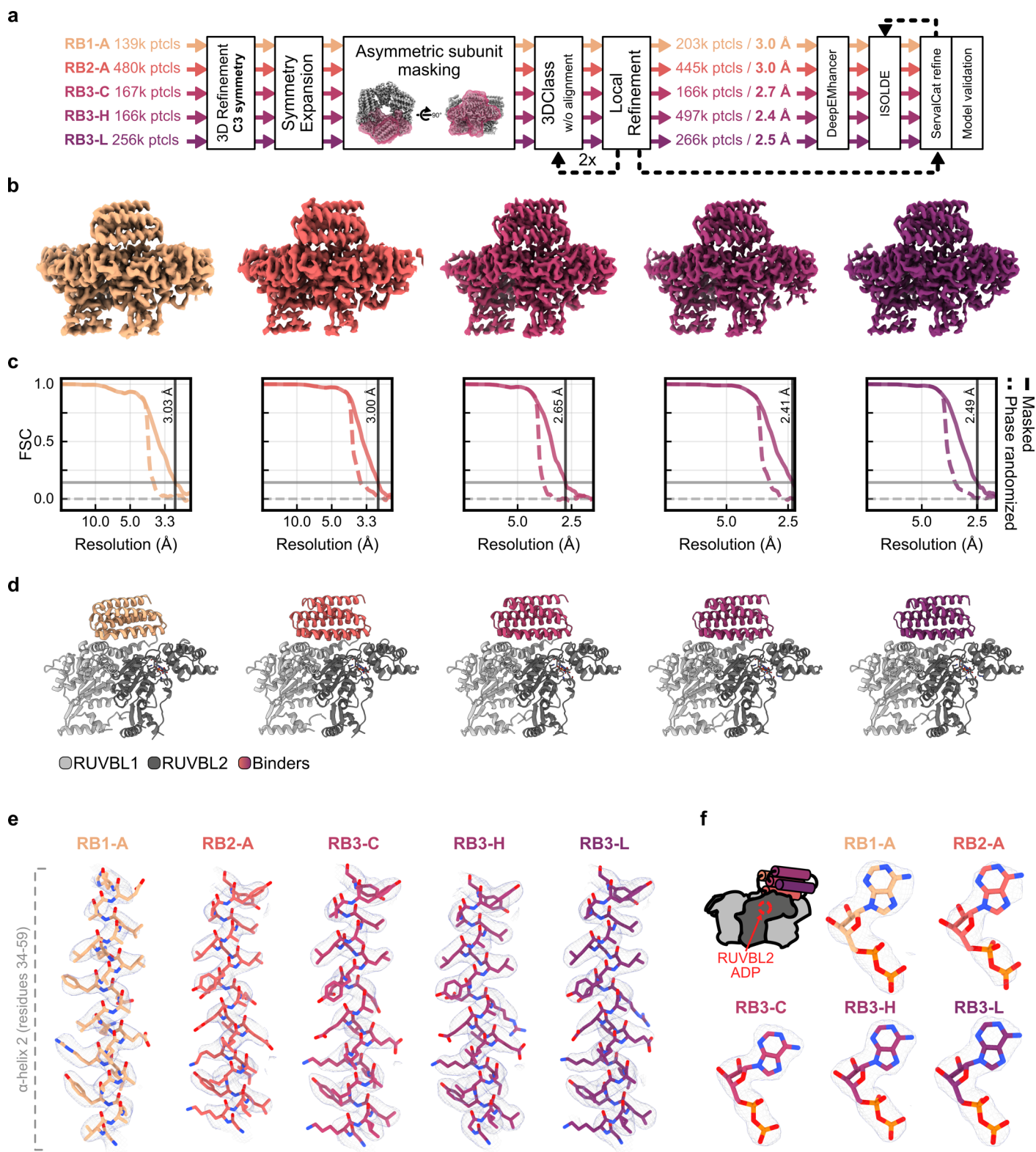

**Supplementary Figure S5. CryoEM structure of the asymmetric unit of RUVBL1/2-RB.** **a**, 3D reconstruction and model building pipeline followed for the asymmetric unit of each RUVBL1/2-RB complex. All resolution values displayed are thresholded at FSC = 0.143. **b**, Density surface representation of the raw, unsharpened reconstruction of the asymmetric unit of RUVBL1/2-RB complexes. **c**, Gold-standard FSC plots of each of the final 3D reconstructions. X-axes are plotted in reciprocal scale (1/x), but x-axis labels correspond to real-space frequency values. Horizontal grey lines indicate FSC = 0 (dashed) and FSC = 0.143 (solid). Vertical black line indicates the frequency value at FSC = 0.143. **d**, Cartoon representation of the RUVBL1/2-RB complex models. **e**, Atomic model (stick representation) fitted into the experimental cryoEM density (mesh surface) for each RB α2 helix. Densities represented here were sharpened using DeepEMhancer for interpretation purposes, but model refinement was performed against gold-standard refined, unsharpened maps. **f**, Same as panel e for the ADP densities within RUVBL2.

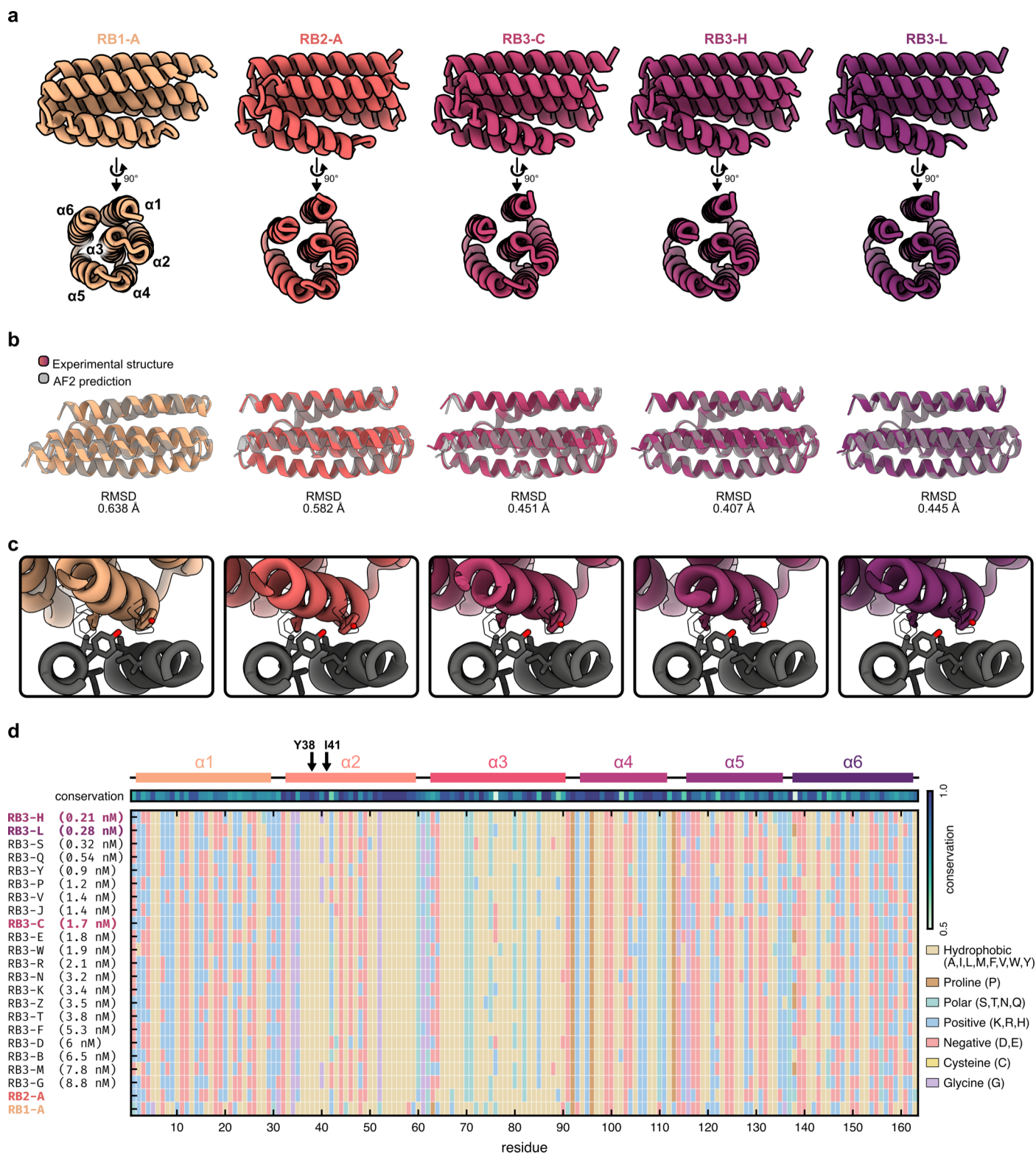

**Supplementary Figure S6. Retrospective analysis of AF2 predictions.** **a**, Cartoon representation of the experimental structures of RBs, two orthogonal views shown. **b**, Structural alignments of each RB experimental structure (colored) and their respective AF2 predictions (grey). The RMSD values were calculated by all-atom structural alignment. **c**, Zoom into Site 1 interaction of RB1-A, RB2-A, RB3-C, -H and -L. Side chains are displayed in stick representation. Side chains of RPAP3 F630 and M626 are represented as black outlines. **d**, Sequence alignment of all RB3s characterized, RB1-A and RB2-A, sorted in descending affinity order.

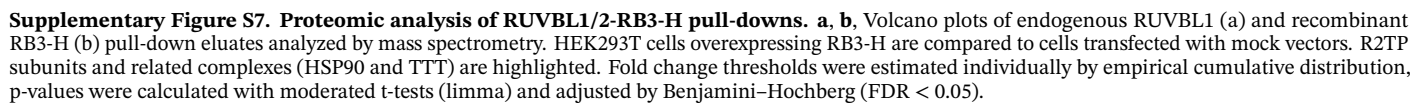

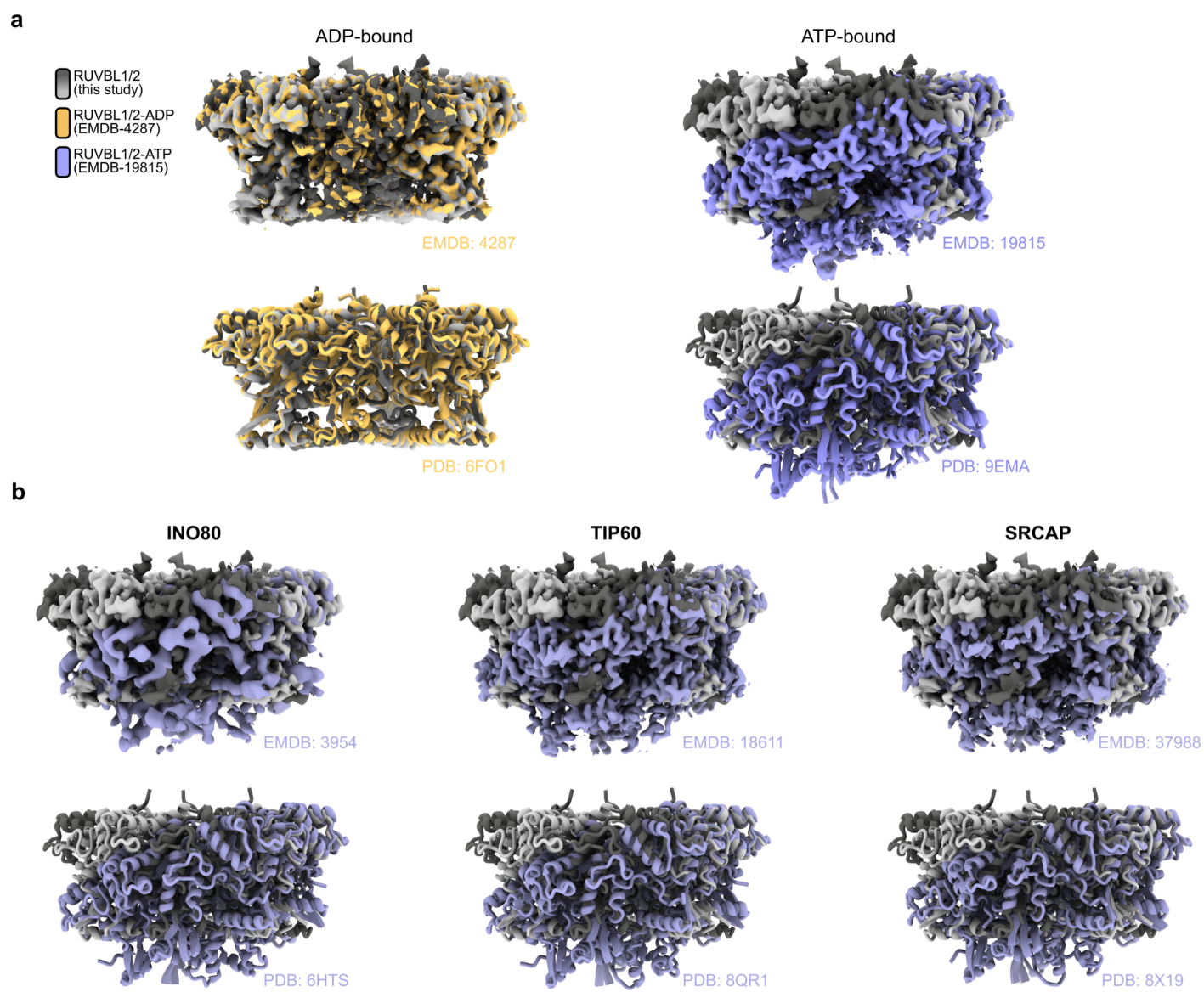

**Supplementary Figure S8. Comparative analysis of RUVBL1/2 conformation upon RB interaction.** **a**, RUVBL1/2 hexamers extracted from the RUVBL1/2-RB3-H structure determined in this study, compared with ADP- (left) and ATP-bound (right) conformations. Top: experimental cryoEM maps; bottom: atomic models in cartoon representation. **b**, Same as panel a, but compared to RUVBL1/2 structures extracted from chromatin remodeling complexes, presenting an ATP-bound-like conformation.

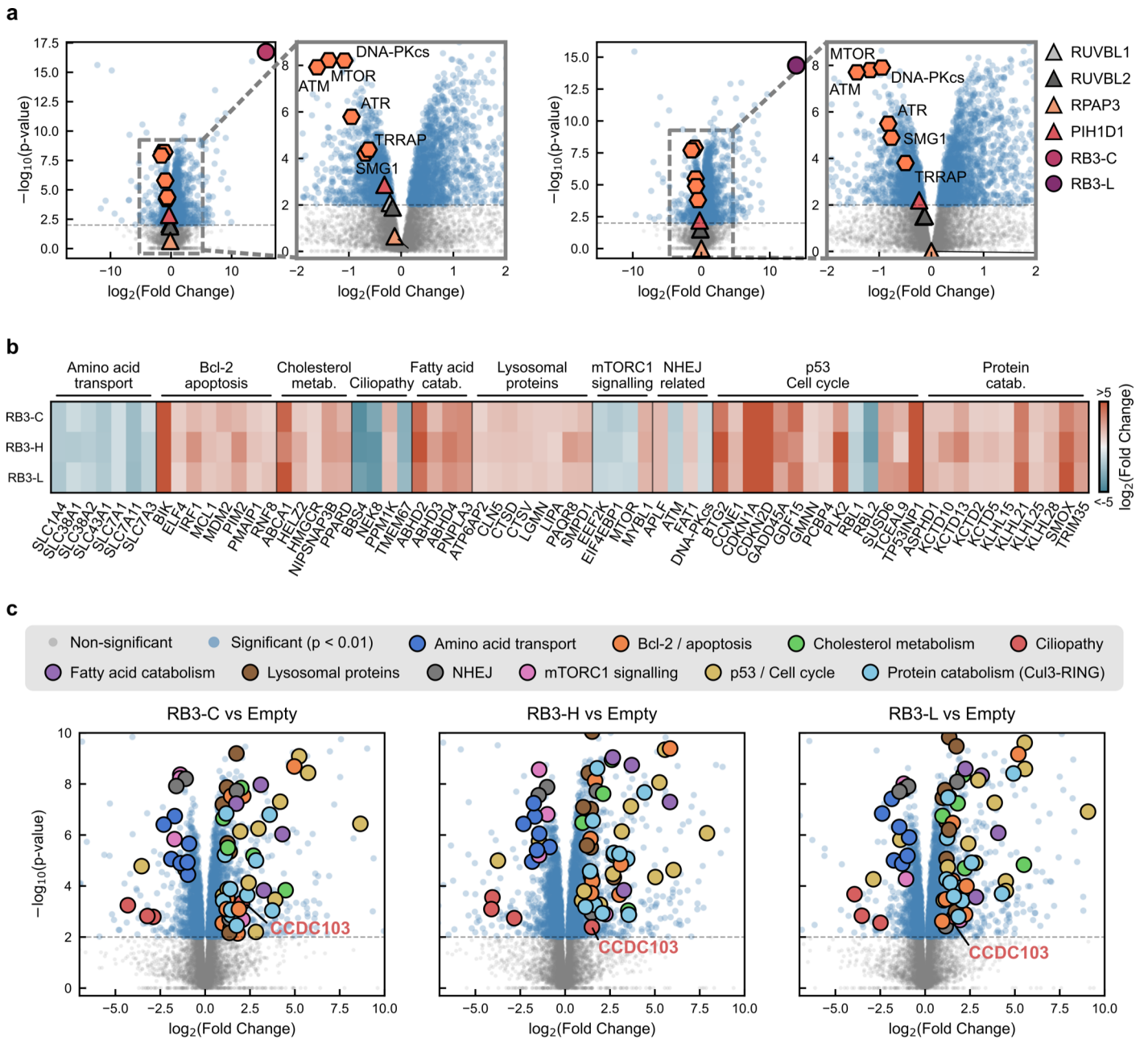

**Supplementary Figure S9. RB3s reduce cellular abundance of all PIKKs.** **a**, Global proteomics analysis volcano plots analogous to Figure 5c for HEK293T cells overexpressing RB3-C (left) and RB3-L (right). $n=3$. **b**, Protein-level heatmap of differential abundance (RB3 overexpression vs control) for RB3-C, RB3-H and RB3-L (rows). Each column is a single protein, cell color encodes $\log_2$(fold change), with color scale centered at 0 and clipped at $\pm 5$. Columns are grouped into functional clusters identified from a STRING-DB run of the proteins showing significant changes across all three RB3s. **c**, Volcano plots for each RB3, as represented in Figure 5c and panel a in this figure. Gray points are non-significant proteins. Larger outlined circles mark members of the STRING-DB clusters also depicted in panel b.

### 2. SUPPLEMENTARY TABLES

Supplementary Table S1. CryoEM and image processing statistics

|  | RUVBL1/2-RB1-A<br>(EMD-XXXX, PDB<br>XXXX) | RUVBL1/2-RB2-A<br>(EMD-XXXX, PDB<br>XXXX) | RUVBL1/2-RB3-C<br>(EMD-XXXX, PDB<br>XXXX) | RUVBL1/2-RB3-H<br>(EMD-XXXX, PDB<br>XXXX) | RUVBL1/2-RB3-L<br>(EMD-XXXX, PDB<br>XXXX) |
| --- | --- | --- | --- | --- | --- |
| **Data collection and processing** |  |  |  |  |  |
| Magnification | 130,000× | 81,000× | 165,000× | 130,000× | 130,000× |
| Voltage (kV) | 300 | 300 | 300 | 300 | 300 |
| Electron exposure (e<sup>-</sup>/Å<sup>2</sup>) | 40.1 | 52.5 | 40.1 | 40.1 | 40.1 |
| Defocus range (μm) | −0.8 to −2.0 | −0.7 to −1.7 | −0.8 to −2.0 | −0.8 to −2.0 | −0.8 to −2.0 |
| Pixel size (Å) | 0.931 | 1.05 | 0.729 | 0.931 | 0.931 |
| Number of movies | 8,383 | 5,105 | 11,863 | 6,772 | 6,036 |
| Symmetry imposed | C1 | C1 | C1 | C1 | C1 |
| Initial particle images (no.) | 379,988 | 781,236 | 551,141 | 1,094,648 | 681,650 |
| Final particle images (no.) | 203,952 (After C3<br>SymExp) | 445,428 (After C3<br>SymExp) | 301,829 (After C3<br>SymExp) | 497,220 (After C3<br>SymExp) | 766,917 (After C3<br>SymExp) |
| Map resolution (FSC=0.143,<br>Å) | 3.1 | 3.0 | 2.7 | 2.4 | 2.5 |
| **Refinement** |  |  |  |  |  |
| Initial model used (PDB<br>code) | 6FO1 + RB AF2<br>prediction | 6FO1 + RB AF2<br>prediction | 6FO1 + RB AF2<br>prediction | 6FO1 + RB AF2<br>prediction | 6FO1 + RB AF2<br>prediction |
| Model resolution (Å) | 3.2 | 3.2 | 2.8 | 2.5 | 2.6 |
| FSC threshold | 0.5 | 0.5 | 0.5 | 0.5 | 0.5 |
| Map sharpening *B* factor<br>(Å<sup>2</sup>) | n/a | n/a | n/a | n/a | n/a |
| **Model composition** |  |  |  |  |  |
| Non-hydrogen atoms | 6,119 | 6,152 | 6,264 | 6,278 | 6,276 |
| Protein residues | 791 | 783 | 796 | 797 | 797 |
| Ligands | ADP (2) | ADP (2) | ADP (2) | ADP (2) | ADP (2) |
| *B* factors (min–max, Å<sup>2</sup>) | 33.30–145.15 | 49.59–148.22 | 12.59–133.36 | 27.22–286.81 | 48.88–150.84 |
| **R.m.s. deviations** |  |  |  |  |  |
| Bond lengths (Å) | 0.0106 | 0.0101 | 0.0101 | 0.0101 | 0.0110 |
| Bond angles (°) | 1.82 | 1.66 | 1.67 | 1.64 | 1.79 |
| **Validation** |  |  |  |  |  |
| MolProbity score | 1.30 | 1.34 | 0.78 | 1.10 | 1.10 |
| Clashscore | 4.27 | 4.16 | 0.94 | 3.05 | 3.13 |
| Poor rotamers (%) | 0.00 | 0.15 | 0.00 | 0.15 | 0.00 |
| **Ramachandran plot** |  |  |  |  |  |
| Favored (%) | 97.55 | 97.27 | 98.72 | 98.22 | 98.47 |
| Allowed (%) | 2.45 | 2.73 | 1.28 | 1.78 | 1.53 |
| Disallowed (%) | 0.00 | 0.00 | 0.00 | 0.00 | 0.00 |
